# Value Representations in the Rodent Orbitofrontal Cortex Drive Learning, not Choice

**DOI:** 10.1101/245720

**Authors:** Kevin J. Miller, Matthew M. Botvinick, Carlos D. Brody

## Abstract

Humans and animals make predictions about the rewards they expect to receive in different situations. In formal models of behavior, these predictions are known as value representations, and they play two very different roles. Firstly, they drive *choice*: the expected values of available options are compared to one another, and the best option is selected. Secondly, they support *learning*: expected values are compared to rewards actually received, and future expectations are updated accordingly. Whether these different functions are mediated by different neural representations remains an open question. Here we employ a recently-developed multi-step task for rats that computationally separates learning from choosing. We investigate the role of value representations in the rodent orbitofrontal cortex, a key structure for value-based cognition. Electrophysiological recordings and optogenetic perturbations indicate that these representations do not directly drive choice. Instead, they signal expected reward information to a learning process elsewhere in the brain that updates choice mechanisms.

## Introduction

Representations of expected value play a critical role in human and animal cognition (Sugrue, Corrado and Newsome, 2005; Lee, Seo and Jung, 2012; Daw and O’Doherty, 2014). A key brain region involved in representing and using expected value information is the orbitofrontal cortex (OFC). An active and unresolved debate in the literature involves theoretical accounts of the OFC that emphasize roles in either choosing (Wallis, 2007; Padoa-Schioppa and Conen, 2017), in learning (Schoenbaum *et al*., 2009; Walton *et al*., 2011; Song, Yang and Wang, 2017), or both (O’Doherty, 2007; Wallis, 2012; Rudebeck and Murray, 2014; Wilson *et al*., 2014; Stalnaker, Cooch and Schoenbaum, 2015). Studies perturbing OFC activity have variously reported behavioral effects consistent with either altered learning (Takahashi *et al*., 2009; Walton *et al*., 2010; McDannald *et al*., 2011; Jones *et al*., 2012; Gardner *et al*., 2019, 2020), or with altered choosing, including conditions involving choosing between multiple possible actions (Murray *et al*., 2015; Ballesta *et al*., 2020; Kuwabara *et al*., 2020) and involving choosing the frequency with which to perform a single action (Jones *et al*., 2012; Gremel and Costa, 2013). Recording studies in many species have revealed neural correlates of expected value in the OFC (Thorpe, Rolls and Maddison, 1983; Schoenbaum, Chiba and Gallagher, 1998; Gottfried, O’Doherty and Dolan, 2003; Padoa-Schioppa and Assad, 2006; Sul *et al*., 2010), but it has not been clear whether these neural correlates of expected value are selective for roles in learning and in choosing. This, along with limitations inherent in neural perturbation studies, has made it difficult to determine whether the OFC plays a role in learning, choosing, or both.

Perturbation studies designed to separate learning from choosing have typically adopted a temporal strategy, attempting to behaviorally separate the time of learning from the time of choosing, and performing neural silencing experiments specifically at one time or the other. Some experiments of this type are consistent with a role for the OFC in learning: silencing the OFC specifically during the learning phase of either of two associative learning procedures (blocking or overexpectation) impairs subsequent behavior on an assay performed with the OFC intact (Takahashi *et al*., 2009; Jones *et al*., 2012). Other experiments of this type are consistent with a role for the OFC in choosing: silencing the OFC specifically during the post-learning assay of either of two associative learning procedures (sensory preconditioning or outcome devaluation) impairs expression of knowledge gained earlier (Jones *et al*., 2012; Gremel and Costa, 2013; Murray *et al*., 2015).

One limitation of studies that attempt to separate learning and choosing temporally is that these are fundamentally cognitive events, internal to the brain, and their timing cannot be fully controlled by an experimenter. For example, it has been suggested that subjects faced with a choice might reconsider information that was presented in the past; cognitively, this might best be interpreted as a learning process happening during a putative choice epoch (Gershman, Markman and Otto, 2014; Lombrozo, 2017; Ludvig *et al*., 2017). Conversely, subjects experiencing a surprising outcome might consider its implications for future choices that they expect to be faced with, and memorize decisions for later use (McDaniel and Einstein, 2007); cognitively, this might best be interpreted as a choice process happening during a putative learning epoch. There is no *a priori* requirement for learning and choosing to be implemented in separate neural circuits. The approach we used here does not depend only on timing, but importantly also uses signals internal to the brain (firing rates, effect of OFC perturbations), to quantify the relative contributions of learning and choosing in the OFC. A second limitation is of previous studies is that they typically address forms of learning that unfold over many sessions, a very different timescale and behavioral regime to the one in which neural representations of value in the OFC are typically studied. Since different neural mechanisms may be at play in trial-by-trial vs. session-by-session learning, it is difficult to confidently interpret the results of neural recording experiments addressing the former in light of neural silencing experiments addressing the latter.

Studies characterizing neural correlates of expected value in the OFC have used a different set of behavioral methods. Typically, they adopt a task design which facilitates the analysis of neural recording by allowing data to be aggregated over many repeated trials. In each trial, the subject is presented with one or more options, chooses among them, and receives a reward associated with the chosen option. Some studies (Thorpe, Rolls and Maddison, 1983; Wallis and Miller, 2003; Sul *et al*., 2010; Costa and Averbeck, 2020) present options that are novel or have repeatedly-changing reward associations in order to elicit trial-by-trial learning, interleaving decision-making with learning by design. Other studies (Tremblay and Schultz, 1999; Padoa-Schioppa and Assad, 2006; Kennerley *et al*., 2009; Rudebeck *et al*., 2013; Blanchard, Hayden and Bromberg-Martin, 2015; Constantinople *et al*., 2019) present options whose reward associations are stable and well-learned, with the intention of isolating decision-making specifically. Although not incentivized by task design, trial-by-trial learning is frequently evident in these tasks as well (Padoa-Schioppa, 2013; Constantinople *et al*., 2019; Lak *et al*., 2020). The fact that learning and choosing are intertwined in these tasks, operating at similar times and over the same set of items, has made it difficult to determine whether OFC value representations are selective for one or the other process, or whether they play a role in both.

Here, we adopt a computational approach that separates learning from choosing, even while using a repeated-trials task that facilitates analysis of neural recordings. This approach is based not only on *when* learning versus choosing happens, but also on *what content* they operate over. We use a multi-step decision task (Daw *et al*., 2011; Miller, Botvinick and Brody, 2017), in which a choice made on a first step is linked probabilistically to an outcome that occurs on a second step, which in turn is linked probabilistically to reward (**Fig. 1a**). Rats adopt a model-based planning strategy that respects this structure (**Fig. 2a**). On the first step of every trial rats make their choice based on values that are computed, not learned (“Compute Choice Port Values” and “Choose a Choice Port” in **Fig. 2a**), while on the second step they learn the values of the outcomes (“Learn Outcome Port Values” in **Fig. 2a**) but are led to those outcomes without making a choice. This task structure, in which choices and outcomes are linked probabilistically rather than having a 1-to-1 relationship, combined with the planning strategy rats use to solve it, provides the critical separation necessary to differentiate learning from choosing.

**Figure 1.**
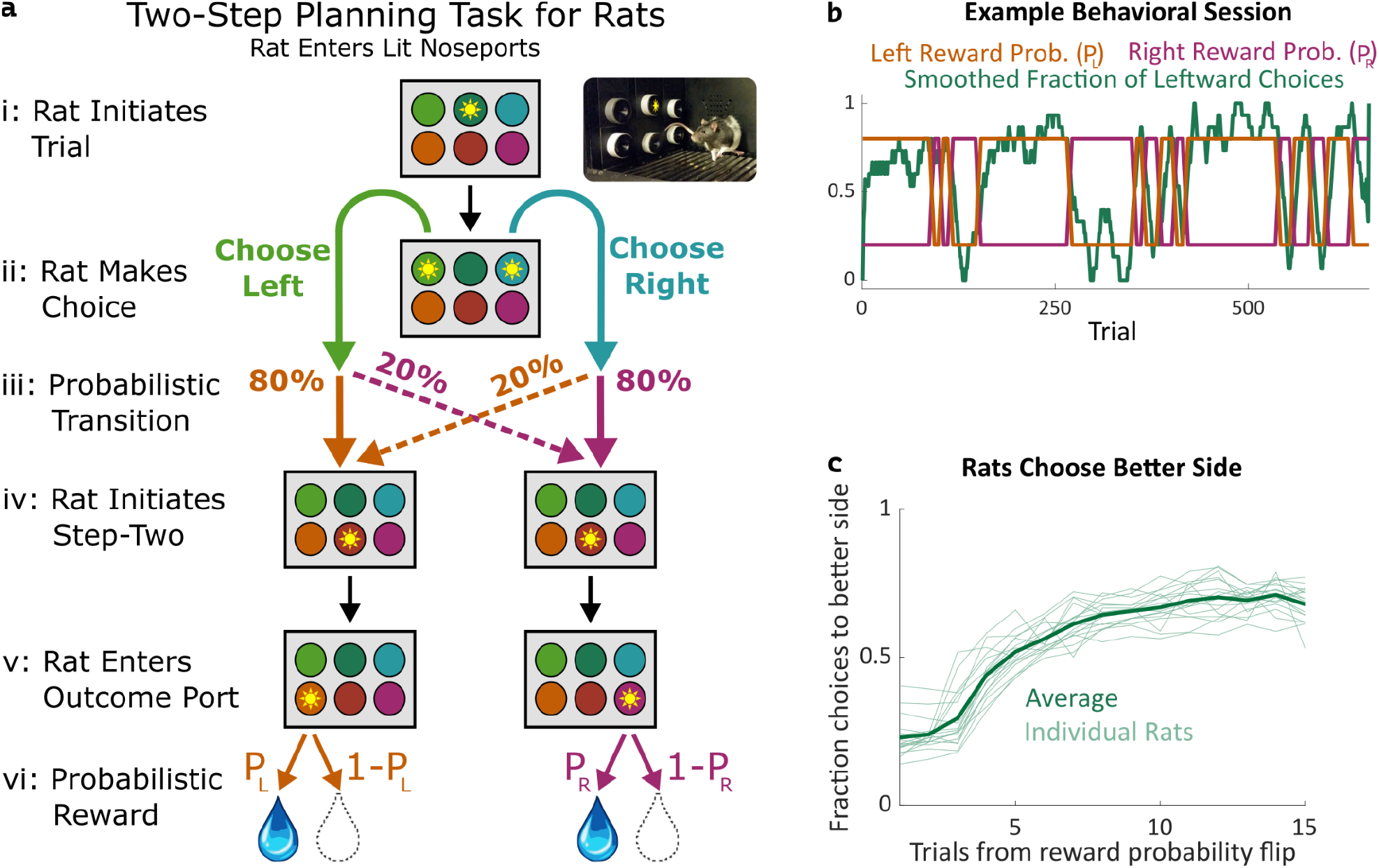
Two-step task for rats. **a:** Rat two-step task. The rat initiates a trial by entering the top center port (i), then chooses to enter one of two choice ports (ii). This leads to a probabilistic transition (iii) to one of two possible paths. In both paths, the rat enters the bottom center port (v), causing one of two outcome ports to illuminate. The rat enters that outcome port (v), and receives a reward (vi). **b:** Example behavioral session. At unpredictable intervals, outcome port reward probabilities flip synchronously between high (80%) and low (20%). The rat adjusts choices accordingly. **c**: The fraction of trials on which each rat selected the choice port whose common (80%) transition led to the outcome port with the currently higher reward probability, as a function of the number of trials that have elapsed since the last reward probability flip.

**Figure 2:**
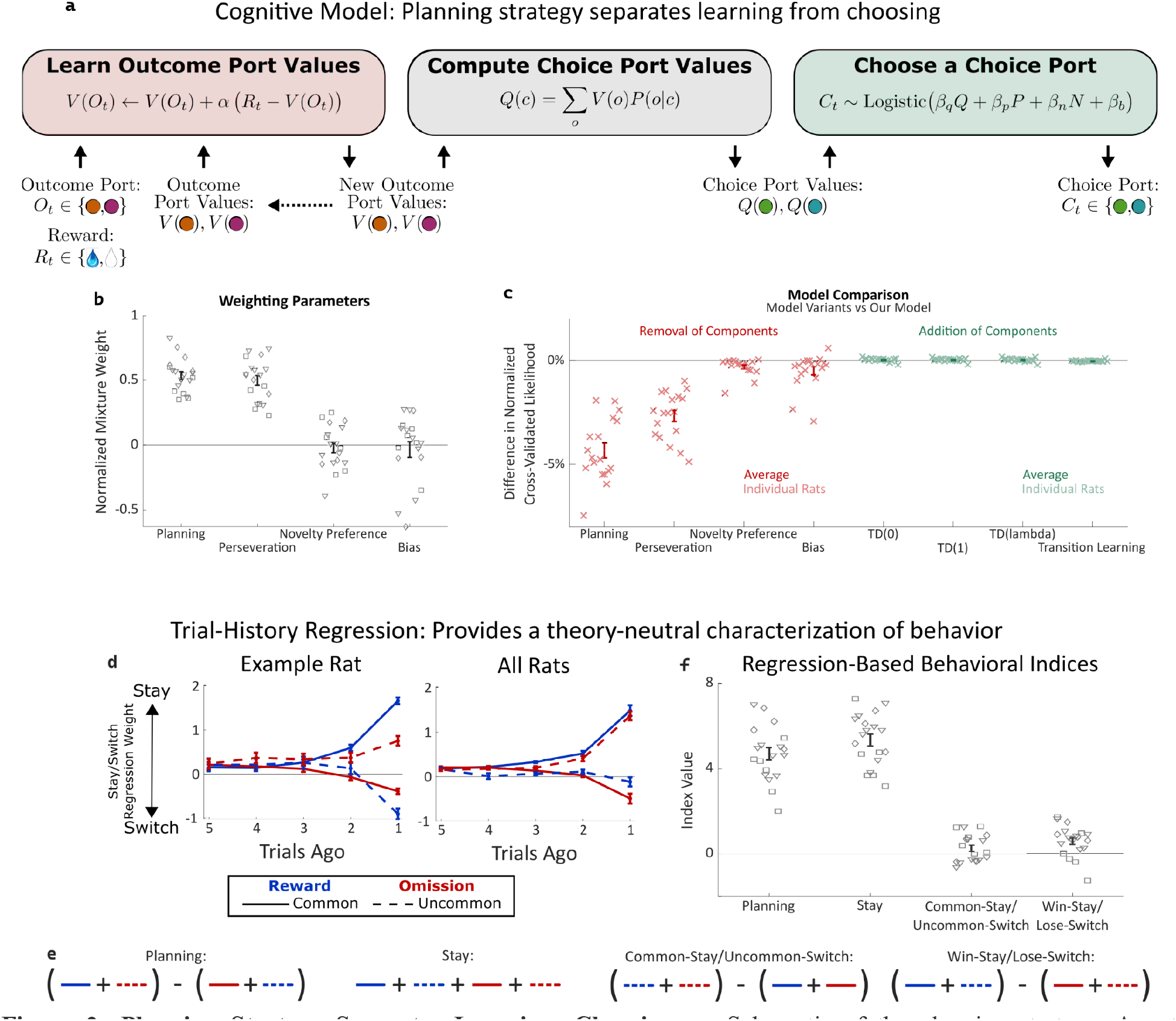
Planning Strategy Separates Learning, Choosing. **a:** Schematic of the planning strategy. Agent maintains value estimates (*V*) for each outcome port, based on a history of recent rewards at that port; as well as value estimates (*Q*) for each choice port, which are computed on each trial based on the outcome values and the world model (*P(o*|*c)*). Choices are drawn probabilistically, based on a weighted combination of these values and of the influence of three other behavioral patterns: perseveration, novelty preference, and bias (see Methods for details). **b**: Mixture weights of the different components of the cognitive model fit to rats’ behavioral data. shown for electrophysiology rats (n=6, squares), optogenetics rats (n=9, triangles), and sham optogenetics rats (n=4, diamonds). **c:** Change in quality of model fit resulting from removing components from the model (red) or adding additional components (green). **d**: Fit weights of the trial-history regression for an example rat (left) and averaged over all rats (right). **e:** Definitions of the four behavioral indices in terms of the fit stay/switch regression weights. The planning index for a particular rat is defined as the sum of that rat’s common-reward and the uncommon-omission weights, minus the sum of its common-omission and uncommon-reward weights. The “stay” index is defined as the sum of all weights. **f:** Values of the four behavioral indices for all rats. The planning and stay indices are large and positive for all rats, while the common-stay/uncommon-switch and win-stay/lose-switch indices are smaller and inconsistent in sign.

We asked whether OFC neuron firing rates were correlated with the expected values of the items being learned about, the values being chosen between, or both. We find that neurons in the OFC correlate significantly with the expected values of the items being learned about, but only weakly with the expected values of the items being chosen between. This indicates that, within a repeated-trials task, neural representations in the OFC carry value information that is largely selective for a role in learning, but only weakly carry value information selective for a role in choice. To causally probe whether OFC plays a role in learning, in choosing, or in both processes, we transiently silenced OFC activity, and found that the pattern of behavior induced by this silencing was reproduced in our computational model only when we disrupted only the role of expected value in learning. Disrupting the role of value in choice did not reproduce this effect. In this model, learning could still occur, but was impaired due to one of its key input signals, expected value, being disrupted.

These results suggest that, even when choosing and learning are commingled on the same trial-by-trial timescale, value representations in the rodent OFC do not drive choices directly, but instead support learning. Moreover, we identify a specific computational role for OFC within learning: it supplies information about expected outcomes to a separate learning process, elsewhere in the brain, that updates value estimates, which can then in turn be used to drive future behavior (Schoenbaum *et al*., 2011).

## Results

### Planning Strategy in the Two-Step Task Separates Choosing and Learning

We trained rats to perform a multistep decision-making task in which they adopt a strategy of model-based planning (Miller, Botvinick and Brody, 2017). The structure of the task was as follows: The rat initiated each trial by poking its nose into a neutral center port, and then selected one of two choice ports (**Fig. 1a i,ii**). One choice caused a left outcome port to become available with probability 80% (“common” transition), and a right outcome port to become available with probability 20% (“uncommon” transition), while the opposite choice reversed these probabilities (**Fig. 1a iii**). These transition probabilities were fixed for each rat, but counterbalanced across rats. Following the initial choice, an auditory cue informed the rat which of the two outcome ports had in fact become available on that trial and, after poking into a second neutral center port, the available outcome port was further indicated by a light (**Fig. 1a iv,v**). The rat was required to poke into the available outcome port (no choice in this step), where it received a water reward with some probability (**Fig. 1a v,vi**). The reward probability at each outcome port on each trial was either 80% or 20%, and these probabilities reversed at unpredictable intervals (**Fig. 1b**). Earning a high reward rate on this task requires learning which outcome port currently has the higher reward probability, and choosing the choice port that is more likely to lead to that outcome port. Rats did this successfully, switching their average choices to the appropriate choice port within several trials of a reward probability reversal (**Fig 1b, c**).

A high reward rate on this task can be achieved by multiple cognitive strategies (Daw *et al*., 2011; Kool, Cushman and Gershman, 2016). In previous work (Miller, Botvinick and Brody, 2017), we showed that rats solve the task using a particular strategy termed “model-based planning” (Dolan and Dayan, 2013; Miller and Venditto, 2021), and presented a cognitive model implementing this strategy (**Fig 2a**). Model-based planning in our task involves separate representations of the expected value associated with the outcome ports and of the expected value associated with the choice ports, and very different computational roles for each type of value representation. Outcome port values, (labeled *V(o)* in our model), represent an estimate of the reward that can be expected following a visit to the corresponding outcome port (*o*). They are updated incrementally by a learning process that compares this expectation to the reward actually received on each trial (labeled *R*_*t*_, **Fig. 2a**, left: “Learn Outcome Port Values”). We used a symmetric update for the value of the outcome port that was not visited (see Methods). Choice port values (labeled *Q(c)* in our model) represent an estimate of the reward that can be expected following a visit to the corresponding choice port (*c*). They are computed based on the known transition probabilities between first-step choice and second-step outcome (the “world model”, *P(o*|*c)*; **Fig. 2a**, center: “Compute Choice Port Values”). These choice port values are then used to determine the next choice (**Fig. 2a**, right: “Choose a Choice Port”). The values of the choice ports therefore drive choice directly, while the learned values of the outcome ports support choice only indirectly, by directly supporting learning.

As in our previous study (Miller, Botvinick and Brody, 2017), rat behavior was well-described by a cognitive model combining this model-based planning strategy with a mixture of three additional components. The first of these is “perseveration”, which reflects a tendency to repeat past choices, regardless of their outcomes (Akaishi *et al*., 2014; Miller, Shenhav and Ludvig, 2019). The second, which we term “novelty preference”, reflects a tendency to repeat (or to switch away from) choices that lead to an uncommon transition, regardless of whether or not they are rewarded. The third is a constant side bias, reflecting an overall tendency to prefer either the right or the left choice port. Each of these components is associated with a weighting parameter (*β*), reflecting the strength of its influence on the decision between the left and right choice port on each trial (**Fig 2a**, right: “Choose a choice port”). Fitting these parameters to the dataset for each rat, we find that the planning and perseverative components earn weights that are large and positive (**Fig 2b**), while the novelty preference and bias components earn weights that are generally smaller and differ in sign among rats (**Fig 2b**). The planning and perseveration components are each associated with a learning rate parameter (*α*), which reflects the relative influence of trials in the recent past vs the more distant past. Learning rates for the planning component were consistently larger than those for the perseverative component (**Fig 2-S1**).

We validate that our cognitive model provides a good description of rat behavior in two different ways. The first way is quantitative model comparison: we compute a quality-of-fit score for the model using cross-validated likelihood, and compare this score between our model and various alternatives (**Fig 2c**). Alternative models which are missing any of the four components perform substantially worse (**Fig 2c**, red points), while alternative models adding various additional components do not perform substantially better (**Fig 2c**, green points). Among these additional components we tested were several model-free reinforcement learning strategies, which have been reported to contribute to behavior on similar multi-step tasks (Daw *et al*., 2011; Hasz and David Redish, 2018; Dezfouli and Balleine, 2019; Groman *et al*., 2019; Akam *et al*., 2020; Miranda *et al*., 2020). That adding them to our model does not improve quality of fit suggests that model-free reinforcement learning does not contribute meaningfully to rat behavior in our task. This is important for our analysis because these strategies involve value representations of their own – that they are not part of our model means that model-based value representations are the only representations of expected reward that are present.

Our second way of validating the cognitive model makes use of a trial-history regression analysis (Lau and Glimcher, 2005), which provides a theory-neutral way of characterizing the patterns present in a behavioral dataset. This analysis fits separate weights for each of the four possible outcome types (common-rewarded, common-omission, uncommon-rewarded, uncommon-omission). These weights will be positive if the rat tends to repeat choices that lead to that outcome, and negative if the rat tends to switch away from such choices. Behavioral datasets produced by different strategies will have different patterns of weights (Miller, Brody and Botvinick, 2016). For example, a model-based planning strategy will show more positive weights for common-reward than for uncommon-reward (since a rewarding outcome port visited after a common transition is likely to be reached again by repeating the choice, while one reached after an uncommon transition is more likely to be reached by switching to the other choice port instead), as well as more negative weights for common-omission than for uncommon-omission (since a unrewarding outcome port visited after a common transition is likely to be avoided by switching the choice, while one reached after an uncommon transition is more likely to be avoided by repeating the choice). As in our previous study, rats trained for the present study universally show this qualitative pattern, and the quantitative patterns present in their weights are well-matched by fits of the cognitive model (example rat: **Fig 2d**; all rats: **Fig 2-S2**). To summarize the pattern and compare it to others in behavior, we define a “planning index” as the linear combination of weights consistent with the planning strategy, as well as a “stay” index, a “common-stay/uncommon-switch” index, and a “win-stay/lose-switch” index quantifying other patterns (**Fig 2e**). All rats showed large values of the planning index and the stay index, and much smaller values of the common-stay/uncommon-switch and win-stay/lose-switch indices (**Fig 2f**).

This cognitive model allows us to probe the role of the OFC in two key ways. First, the model can be run using choices and rewards actually experienced by the rat to provide trial-by-trial timecourses for the expected values of the choice ports (*Q*) and outcome ports (*V*). We will use these timecourses (Daw, 2011) as estimates of the value placed by the rat on the various ports, and look for correlates of them in the activity of OFC neurons. Second, the model can be altered selectively and run to generate synthetic behavioral datasets, providing predictions about the specific behavioral effects of particular cognitive impairments. We will selectively disrupt choice port value information (which directly drives choice) or outcome port value information (which participates in learning) within the model on a random subset of trials. We will compare these synthetic datasets to real behavioral datasets from rats in which we silence neural activity in the OFC.

### Neural Activity in Orbitofrontal Cortex Codes Expected Value of Outcomes

We implanted a recording array into the left OFC (**Fig 3d, Fig 3-1**) of each of six rats, and performed electrophysiological recordings during 51 behavioral sessions. This dataset yielded 477 activity clusters, including both single-unit and multi-unit recordings. We characterized whether the activity of each unit encoded three different types of value information, which we estimated trial-by-trial using fits of the cognitive model (Daw, 2011). The first type was the difference in expected value between the left and the right choice ports, which we term “choice value difference”. Representations of this kind would be consistent with a role in driving choice. The second was the expected value of the choice port that the rat actually selected on that trial, termed “chosen value”. This signal has also been proposed to play a role in the choice process (Rustichini and Padoa-Schioppa, 2015). The last was the expected value of the outcome port visited by the rat on that trial, which we term “outcome value”. Representations of outcome value are predictions about immediate reward probability, and are consistent with a role in learning. We sought to determine whether our neural recording data contained correlates of choice-related value signals, outcome-related value signals, or both.

**Figure 3.**
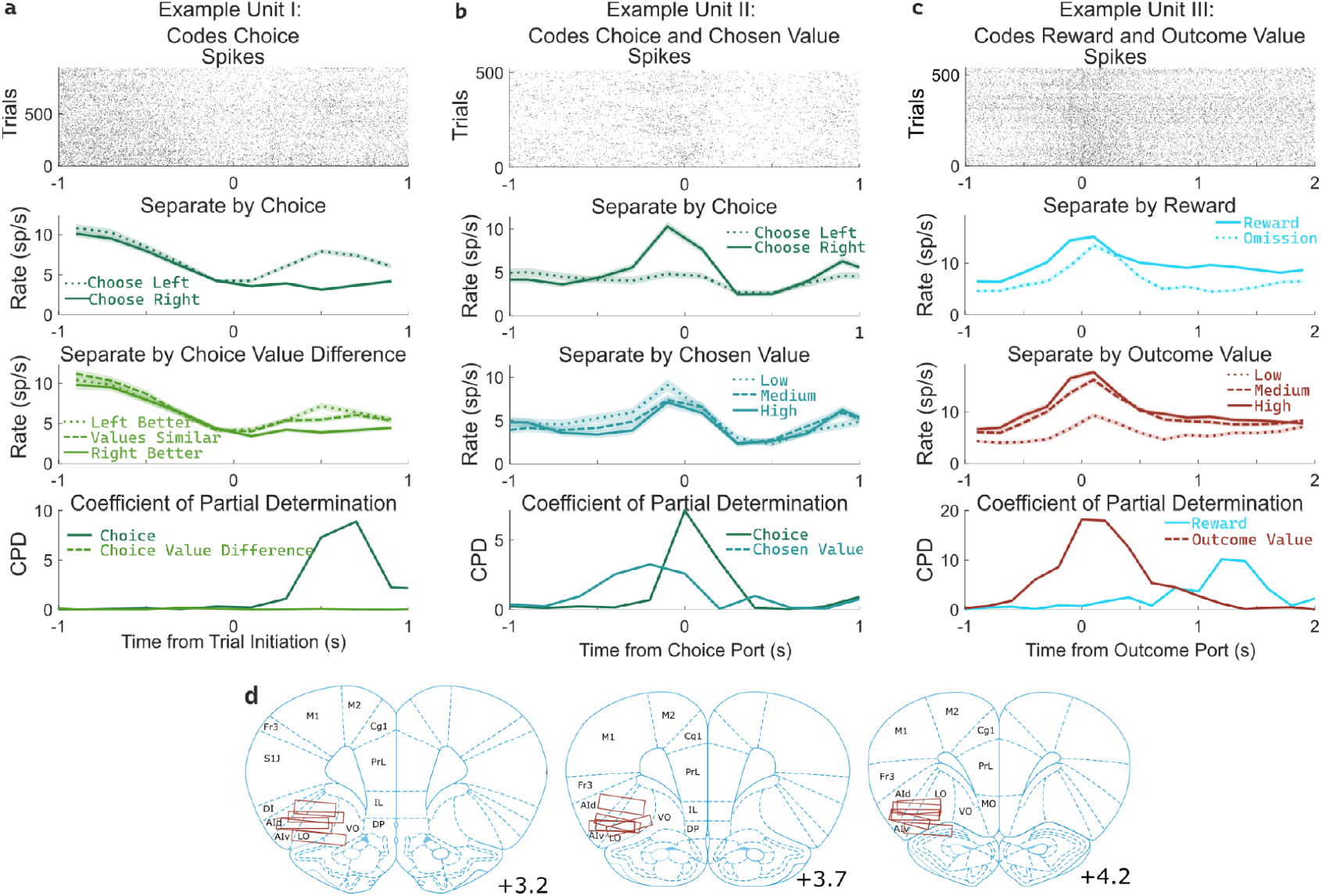
OFC units encode multiple correlated variables. **a**. Example unit whose firing rate differs both with the rat’s choice and with the difference in value between the two possible choices. An analysis using coefficient of partial determination (CPD) reveals choice coding, but no coding of expected value. **b**. Example unit whose firing rate differs both with the rat’s choice and with the expected value of the choice port visited on that trial. CPD analysis reveals coding of both of these variables, though with different timecourses. **c**. Example unit whose firing rate differs both with reward received and with the expected value of the outcome port visited. CPD analysis reveals coding of both of these variables, with different timecourses. **d**. Approximate location of recording electrodes targeting OFC, which in rats is represented by regions LO and AIv (Paxinos and Watson, 2006; Price, 2007; Stalnaker, Cooch and Schoenbaum, 2015), estimated using histology images (Fig3-S1).

Determining whether a unit encodes these quantities requires separating their influence from that of other variables with which they may be correlated. For example, if the firing rate of a unit differs between left-choice and right-choice trials, it will also differ between trials in which the left port had a higher value and those in which the right port had a higher value (**Fig. 3a**). This happens because the rat is more likely to select the choice port that is currently higher-valued. Similarly, if the firing rate of a unit differs between rewarded and unrewarded trials, it will also differ between trials with a high and a low outcome port value (**Fig. 3c**). To quantify coding strength in light of these and other correlations (**Fig. 3-S2**), we used a multiple regression approach. Specifically, we fit a separate regression model to predict the spike counts of each unit in each of several 200ms time bins taken relative to the port entry events, using as regressors the potentially learning-related events from one trial (choice port, outcome port, reward, and their interactions, as well as outcome value) as well as the potentially choosing-related variables from the next trial (choice port, choice value difference, and chosen value). We then computed for each regressor the coefficient of partial determination (CPD, also known as ‘partial r-squared’; Cai, Kim and Lee, 2011; Kennerley, Behrens and Wallis, 2011), which quantifies the percentage of remaining variance explained by that regressor, once the influence of all other regressors has been accounted for (**Fig. 3abc**, bottom row).

Coefficients of partial determination can be computed for a particular fit (one unit in one time bin), or for a collection of fits (aggregating variance over units, bins, or both). First, we considered coding in individual clusters (single- or multi-unit), aggregating data from time bins within a one-second window around the time of each nose port entry. We assess significance by constructing for each unit a set of null datasets by circularly permuting the trial labels, and comparing the CPDs in the true dataset to those in the null datasets. Using this method, we found that a large fraction of units significantly modulated their firing rate according to outcome-value, with the largest fraction doing so at outcome port entry (170/477, 36%; permutation test at p<0.01). In contrast, a relatively small fraction of units modulated their firing rate according to choice-value-difference (largest at outcome port entry: 34/477, 7%), or to chosen-value (largest at choice port entry: 61/477, 13%). Furthermore, the magnitude of CPD was larger for outcome-value than for the other value regressors. Considering the port entry event with the strongest coding for each regressor, the mean cluster had CPD for outcome-value 3.5x larger than for choice-value-difference (p=10^−24^, sign rank test; median unit 1.6x larger; **Fig. 4a**, note logarithmic axes), and 3.7x larger than for chosen-value (p=10^−14^; median unit 1.7x; **Fig. 4a**). Considering each 200ms time bin separately, we find the fraction of units encoding outcome value rises shortly before step two initiation and peaks sharply around the time of outcome port entry, the fraction coding choice value difference peaks at outcome port entry as well, while the fraction coding chosen value peaks at the time of choice port entry (**Fig 4-S1**). Next, we considered coding at the population level, computing CPD over all clusters for each time bin and subtracting the average CPD from the null datasets (**Fig 4b**). We found robust population coding of outcome-value, beginning at the time of entry into the bottom-center port, and peaking shortly after entry into the outcome port at 0.82%. In contrast, population coding of choice-value-difference and chosen-value was low in all time bins, reaching a maximum of only 0.29%. Population coding in the OFC was also present for other regressors in our model, especially for reward, choice, and outcome port (**Fig 4c**). Similar results were found when considering single- and multi-unit clusters separately (**Fig 4-S2, Fig 4-S3**), as well as when separating the units recorded from individual rats (**Fig 4-S4, Fig4-S5**). These results were robust to replacing the choice-related value regressors with analogs that consider the full decision variable, including contributions from perseveration and novelty preference as well as expected value (**Fig 4-S6**). They were also robust to removing a regressor (interaction of reward and outcome port) that was particularly highly correlated with choice value difference (**Fig 4-S6**). Across all of these variants a clear pattern was present: neural activity in OFC encodes the expected value of the visited outcome port more strongly than it encodes either type of value information about the choice ports. In computational models (**Fig 2a**), this type of value information plays a role in supporting learning, but does not play a direct role in choice. Our neural recording results therefore suggest that the OFC may play a role in supporting learning, but cast doubt on the idea that they play a strong role in driving choice directly.

**Figure 4.**
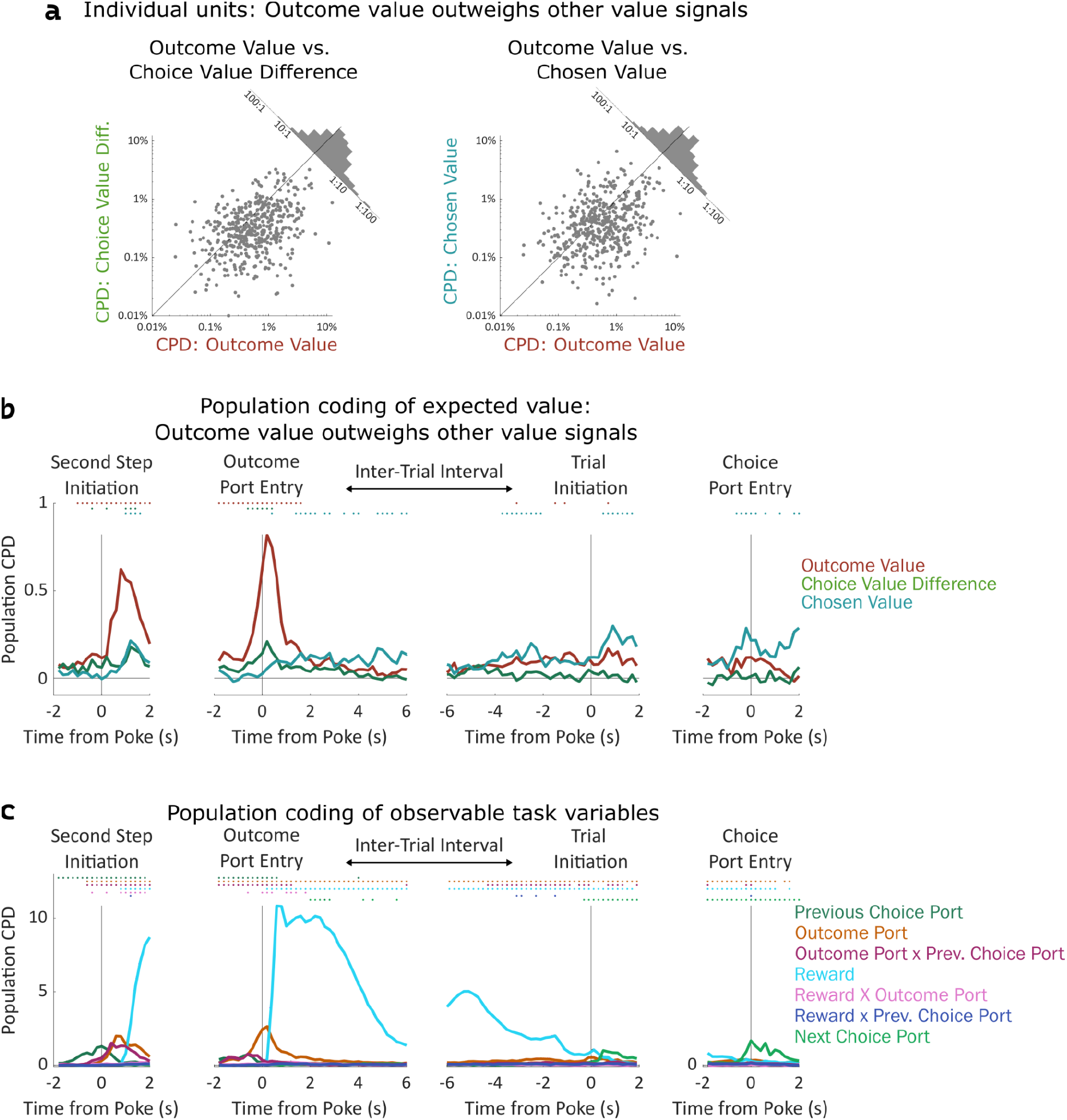
Coding of expected value of outcomes outweighs coding of expected value of choices. **a**. Left: Scatterplot showing CPD for each unit (n=477) for the outcome-value regressor against CPD for the choice-value-difference regressor, both computed in a one-second window centered on entry into the outcome port. Right: Scatterplot showing CPD for outcome-value, computed at outcome port entry, against CPD for the chosen-value regressor, computed at choice port entry. **b**. Timecourse of population CPD for the three expected value regressors. We have subtracted from each CPD the mean CPD found in permuted datasets. **c**. Timecourse of population CPD for the remaining regressors in the model, which reflect observable variables and interactions between them.

### Inactivations of OFC Impair Update Process, not Choice Process

To assess the causal role of the OFC’s value signals, we silenced neural activity using the optogenetic construct halorhodopsin (eNpHR3.0; **Fig 5b, Fig 5-S1**) during either the *outcome period* (beginning at entry into the outcome port and lasting until the end of reward consumption), the *choice period* (beginning at the end of reward consumption and lasting until entry into the choice port on the subsequent trial), or *both periods* (**Fig. 5a, Fig 5-S2**) in an experimental group of nine rats (577 total sessions). Previous work (Miller, Botvinick and Brody, 2017) had shown that whole-session pharmacological silencing of the OFC specifically attenuates the “planning index” (**Fig. 2e**, see Methods), which quantifies the extent to which the rats’ choices are modulated by past trials’ outcomes in a way consistent with planning. Here, we found that optogenetic inactivation spanning both the outcome and the reward period reproduced this effect, decreasing the planning index on the subsequent trial (p=0.007, t-test with n=9 rats; **Fig 5c**). Inactivation during the outcome period alone also decreased the planning index on the subsequent trial (p=0.0007, **Fig 5c**), but inactivation during the choice period alone did not (p=0.64). Comparing the strength of the effect across time periods, we found that both reward-period and both-periods inactivation produced effects that were similar to one another (p=0.5), but greater than the effect of choice-period inactivation (p=0.007, p=0.02). We repeated the experiment in a control group of four rats (109 total sessions) that were implanted with optical fibers but did not express halorhodopsin. This sham inactivation produced no significant effect on the planning index for any time period (all p>0.15, t-test with n=4 rats; **Fig 5c**, grey diamonds), and experimental and control rats differed in the effects of outcome-period and both-period inactivation (p=0.02, p=0.02, two-sample t-tests). As in our previous study, inactivation of OFC did not significantly affect other regression-based behavioral indices (**Fig 5-S3**). Together, these results indicate that silencing the OFC at the time of the outcome (and therefore the time of peak outcome-value coding, **Fig 4b**) is sufficient to disrupt planning behavior.

**Figure 5.**
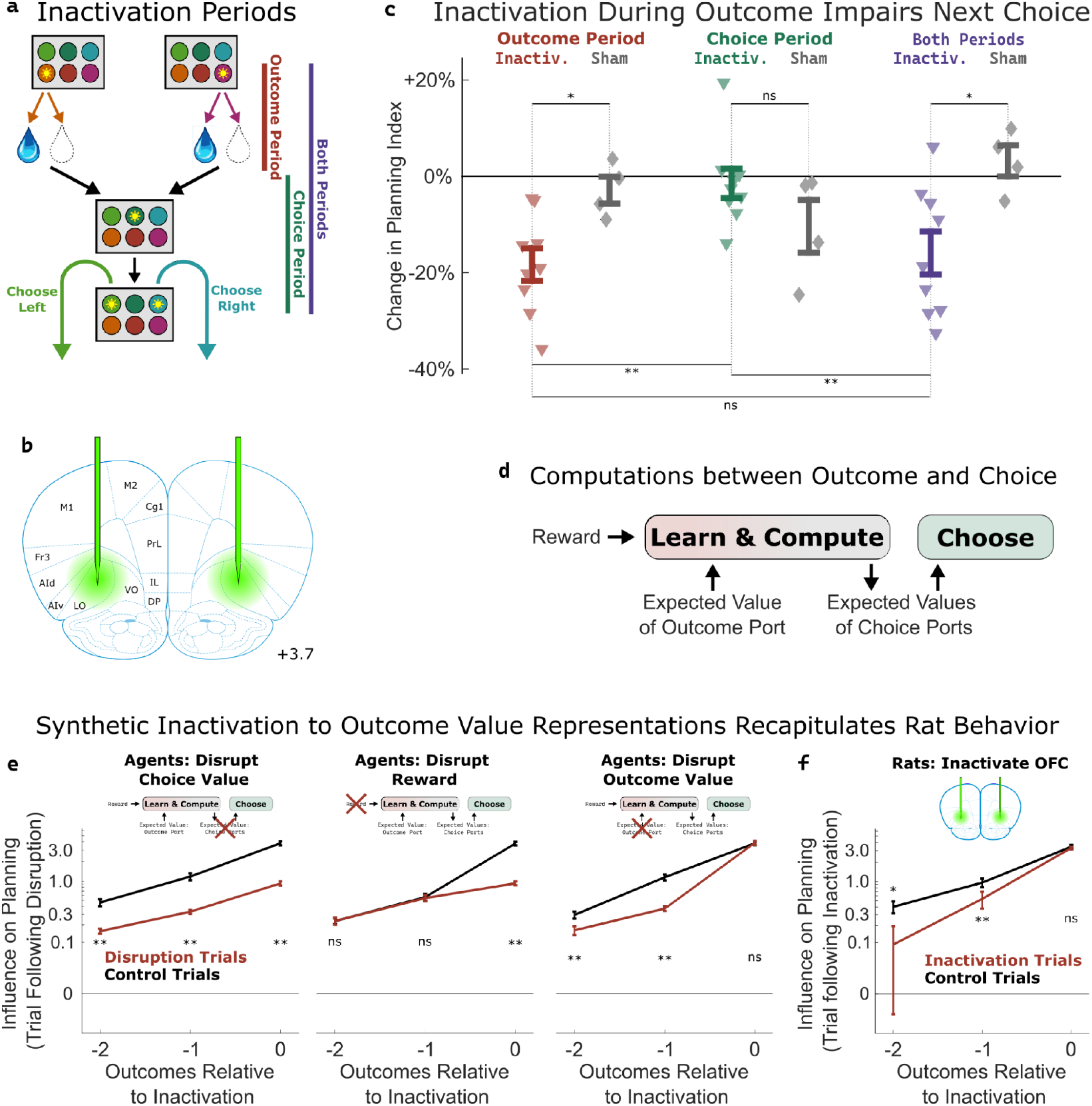
Inactivation of OFC attenuates influence of outcome values. **a:** Three time periods of inactivation. Outcome-period inactivation began when the rat entered the outcome port, and continued until the rat exited the port, or for a minimum of two seconds. Choice-period inactivation began after this outcome period, and continued until the rat entered the choice port on the next trial, or for a maximum of 15 seconds. Both-period inactivation encompassed both of these periods. **b:** Target location for optical fiber implants. See Fig5-S1 for estimated actual locations in individual rats. Coronal section modified from Paxinos and Watson (2006). **c:** Effects of inactivation on the planning index on the subsequent trial for experimental rats (n=9, colored triangles) and sham-inactivation rats (n=4, grey diamonds). Bars indicate standard errors across rats. **d:** Simplified schematic of the representations and computations that take place in our software agent between the delivery of the outcome on one trial and the choice on the next. Compare to Fig 2a. **e:** Analysis of synthetic datasets created by disrupting different representations within the software agent on a subset of trials. Each panel shows the contribution to the planning index of trial outcomes at different lags on choices, both on control trials (black) and on trials following disruption of a representation (red). Bars indicate standard error across simulated rats (see Methods) **f**: Same analysis as in c, applied to data from optogenetic inactivation of the OFC during the outcome period.

To help understand which aspect of the behavior was affected by silencing the OFC, we used our cognitive model (**Fig 2a**) to perform three different types of synthetic inactivation experiments. Each of these produced a distinct pattern of behavior, most clearly visible when we separately computed the contribution to the planning index of the past three trials’ outcomes (**Fig. 5e**). To simulate an effect of inactivation on the choice process, we decreased the value of the planning agent’s weighting parameter (*β*_*q*_; **Fig 2a**, “choose a choice port”) on synthetic inactivation trials, effectively reducing the influence of choice port value representations on choice. This resulted in choices that were more noisy in general, affecting the influence of all past outcomes (**Fig 5e**, left). To simulate effects of inactivation on the learning process, we reparameterized the agent’s learning equation (**Fig 2a**, “update outcome port values”) to facilitate trial-by-trial alterations of learning parameters:

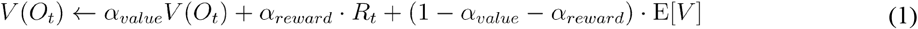

where *α*_*value*_ and *α*_*reward*_ are now separate learning rate parameters, and E[*V*] represents the expected reward of a random-choice policy. We made a corresponding change to the symmetric update for the outcome port that was not visited (see Methods). To simulate an effect of inactivation on the role of reward in learning, we decreased *α*_*reward*_ on synthetic inactivation trials. This resulted in a change specifically to the influence of the outcome during which we performed the synthetic inactivation on future choice (**Fig 5e**, middle). To simulate an effect of inactivation on the role of value in learning, we decreased *α*_*value*_ on synthetic inactivation trials. This resulted in a change to the influence of outcomes from *previous* trials (**Fig 5e** right) but not of the trial during which synthetic inactivation was actually performed. This is because value acts as a summarized memory of previous trials’ outcomes, and attenuating it affects the influence of all of these.

We performed the same analysis on the inactivation data from our rats **(Fig. 5f**), and found that silencing OFC during the outcome period on a particular trial did not affect the influence of that trial’s outcome on the upcoming choice (p=0.2, paired t-test with n=9 rats), but that it did affect the influence of the previous two trials’ outcomes (p=0.004, p=0.02). This pattern was consistent with the synthetic dataset in which outcome value representations had been attenuated, but not the other synthetic datasets (compare **Fig. 5f** to **Fig. 5e**). We conclude that silencing the OFC in our task predominantly impairs the use of outcome port value information, needed for updating value expectations, but has little or no effect on the use of choice port value information, needed for driving choice.

## Discussion

In formal models of value-based cognition, value representations can play two very different computational roles. First, they can drive choosing, as expected values of different available options are compared to one another, and the best is selected. Secondly, they can drive learning, as the expected value of an outcome is compared to the reward actually received, and future expectations are updated. Value representations in the OFC have been reported in many tasks and species, but it is still unclear whether they drive one process, the other process, or both. The rat two-step task gave us the opportunity to separate the two roles, both in terms of coding in neural activity and in terms of the behavioral impact of silencing that activity. Using this approach, we find weak (though significant) representation of values associated with the available choices (“choice values”), little or no effect of silencing OFC at the putative time of choice, and effects of silencing inconsistent with impairing choice values in a computational model. Instead, we find strong representation of values associated with immediately impending reward outcomes (“outcome values”), a strong behavioral effect of silencing OFC at the time of those representations, and effects of silencing that are consistent with specifically impairing the use of outcome values for learning. This pattern of results suggests a much stronger role for OFC in learning than in choosing. More specifically, the results suggest that value information in the rodent OFC may not drive choice directly, but instead provide information about expected outcomes to a separate learning process. In the computational model most consistent with our data, OFC inactivation does not prevent learning, but instead impairs one of the critical inputs to the learning process, namely expected outcome value. Our results therefore are consistent with the idea that learning does not take place in the OFC itself, but in some other structure to which OFC supplies this key input.

These results clarify the role of a neural signal that has been reported over a wide variety of tasks and species (O’Doherty, 2007; Wallis, 2012; Rudebeck and Murray, 2014; Stalnaker, Cooch and Schoenbaum, 2015). In decision tasks, several distinct types of value representation have been reported, but the most commonly observed is “chosen value” (Wallis and Miller, 2003; Sul *et al*., 2010; Kennerley, Behrens and Wallis, 2011): the expected reward associated with the option that the subject chose. The role of this signal is not clear, it has been interpreted both as representing a post-decision signal that provides input to a learning process (Schoenbaum *et al*., 2011), but also as an internal component of the choice process itself (Rustichini and Padoa-Schioppa, 2015). These roles are difficult to fully disentangle using classic tasks, in which both learning and choosing operate over the same set of items. Rat behavior in the two-step task separates these possibilities. If “chosen value” relates to the choice process, we would expect to see representations of the expected value of the port that was chosen (not learned about), while if it relates to the learning process, we would expect to see expected value of the port that was learned about (but not chosen). Our data predominantly provide evidence for the latter alternative in rats.

Our results build on prior work attempting to separate learning and choosing in trial-by-trial tasks in primates. One set of studies used a probabilistic reward-learning task (the “three-armed bandit” task), along with detailed trial-by-trial analysis to quantify deficits in “credit assignment”, attributable to an impaired learning process, as well as deficits in decision-making (Walton *et al*., 2010). Studies using this tool reported that both macaques and humans with lesions to the OFC showed impairments in credit assignment, while those with lesions to a neighboring region, ventromedial prefrontal cortex (vmPFC), showed impairments in decision-making (Noonan *et al*., 2010, 2017). A methodological limitation of these studies is that they employed nonselective lesions, which damage both local neurons (from which electrophysiological signals are recorded) as well as fiber tracts connecting other brain regions (from which they typically are not). A subsequent study (Rudebeck *et al*., 2017) used excitotoxic lesions, which do not damage passing fibers, and found that lesions spanning both OFC and vmPFC affected neither credit assignment nor decision making on this task. Instead, the study found that lesions of another neighboring region, ventrolateral prefrontal cortex (vlPFC), caused impairments in credit assignment (Rudebeck *et al*., 2017). Together, these primate results suggest that neural activity in OFC itself is not necessary either for normal choosing or for normal learning on the three-armed bandit task, but that vlPFC may play a specific role in supporting learning (Murray and Rudebeck, 2018). Mapping these findings to the rodent brain is not entirely straightforward. While the distinction in rodents between OFC and vmPFC is relatively clear, the distinction between OFC and vlPFC is much less so (Price, 2007). Anatomically, many of the criteria that motivate the proposed homology between rodent OFC (typically taken to mean areas LO and AIv) and primate OFC (typically taken to mean areas 11l and 13) may apply to parts of primate vlPFC (especially area 12) as well (Price, 2007; Stalnaker, Cooch and Schoenbaum, 2015). Like in OFC, neurons in vlPFC have been shown to represent expected outcomes (Kobayashi, Pinto de Carvalho and Schultz, 2010; Rich and Wallis, 2014), though these signals have not been characterized as intensively as those in primate OFC. A possibility is that primate OFC and vlPFC are specialized for functions that in rodents are both performed by the OFC.

Our results also build on those of previous studies that attempt to separate learning and choosing by separating these functions in time, typically by tens of minutes or longer. Here, however, we investigated a very different timescale in that learning and choosing were interleaved, both occurring on each trial of the two-step task. Our inactivation results are consistent with previous studies suggesting a role for the OFC in learning (Takahashi *et al*., 2009; Jones *et al*., 2012), but in apparent tension with others finding that inactivation impairs performance on a post-learning assay (Jones *et al*., 2012; Gremel and Costa, 2013; Murray *et al*., 2015). One possibility is that the OFC plays different roles for different timescales, or for these different types of behavior. Another possibility is that the OFC plays a common role across timescales and tasks, supporting a process that updates choice mechanisms – in long-timescale tasks this process may take place during either the learning or the probe session, while in our task it necessarily takes place between the outcome of one trial and the choice on the next.

Our results are consistent with the view that the OFC carries expectancy signals (Schoenbaum and Roesch, 2005; Rudebeck and Murray, 2014) indicating which outcomes are expected to follow from the current state. This view is distinct from the “chosen value” view described earlier in that it claims that these signals indicate the particular identities of the expected outcomes (e.g. different types of foods) rather than the abstract “reward” or “common currency” value found in reinforcement learning or neuroeconomic theories. Our experiment does not speak to the difference between these views, because the only reward available in our task is a water droplet of a fixed size. Our “outcome value” correlates might therefore reflect expectations of this reward (the water droplet) in particular, and play a role in updates based on this expectation. In formal models, this might be described as updating estimates of a state transition probability (e.g. from an “at the outcome port” state to a “receive water droplet” state). Alternatively, “outcome value” might abstract over many different possible rewarding outcomes. In formal models, this might be described as updating estimates of the reward function (e.g. in an “at the outcome port” state).

Our results are also in tension with those of some studies using economic choice tasks, in which subjects make decisions between pairs of well-learned stimuli leading to different quantities of differently flavored rewards. A pair of studies, one in mice (Kuwabara *et al*., 2020) and one in primates (Ballesta *et al*., 2020), uses methods that disrupt OFC neural activity on a particular subset of trials, and reports that subjects’ decisions are disrupted specifically on those trials. Recording studies in these tasks, both in primates and in mice (Padoa-Schioppa and Assad, 2006; Kuwabara *et al*., 2020), have reported neural correlates of the values of the individual options that are available. This “offer value” correlate is distinct from the “chosen value” correlate discussed earlier, and is suitable for a role in driving decisions directly. These results have led to the view that OFC value representations, at least in these tasks and species, play a key role in decision-making (Rustichini and

Padoa-Schioppa, 2015; Padoa-Schioppa and Conen, 2017). Our results indicate that in the rat two-step task, OFC does not play this role, since choice-related value correlates are minimal, and inactivation effects are inconsistent with a direct role in choice. One possible resolution to this tension is a difference between tasks. It has been proposed that OFC plays a role specifically in choices between outcomes that offer different “goods” (Padoa-Schioppa, 2011; e.g. different flavors Kuwabara *et al*., 2020). In primate tasks involving decisions between outcomes that differ in other properties – such as magnitude, probability, or delay – OFC value representations are typically more consistent with chosen-value than with offer-value coding (Wallis and Miller, 2003; Roesch and Olson, 2004; Rich and Wallis, 2016). Casting doubt on this goods-based interpretation are a set of recent results from a flavor-based task in rats (Gardner *et al*., 2017, 2020) showing that silencing the OFC does not affect choice. Another possible resolution is a difference between species. Consistent with either of these possibilities are recent results from a choice task in rats in which options differed in probability and magnitude (but not flavor): rat OFC in this task did not carry correlates of offer value, and silencing impaired trial-history effects rather than the influence of offers on choice (Constantinople *et al*., 2019).

A final possibility is that OFC contributes to a model-based “policy update” process that combines known information about the structure of the world with new information, in order to modify an animal’s behavioral strategy (Gershman, Markman and Otto, 2014; Ludvig *et al*., 2017; Mattar and Daw, 2018). Such a process could happen at different times in different tasks. In our task, policy update might happen between the end of one trial and the beginning of the next, and be interpreted as learning. In economic choice tasks, policy update might happen immediately after the options are presented, but before a choice is made, and be interpreted as related to deliberation. Indeed, experiments analyzing response times suggest that such a deliberative process sometimes occurs during economic choice (Krajbich, Armel and Rangel, 2010), and may be related to moment-by-moment patterns of activity in the OFC (Rich and Wallis, 2016).

Consistent with this idea are results from a series of recent studies using a flavor-based economic choice task for rats (Gardner *et al*., 2017). If rats are pre-fed rewards of one flavor immediately before a task session begins, they normally alter their choice preferences during that session; silencing OFC activity impairs this update (Gardner *et al*., 2019). If rats are presented with a novel pairing of flavors that have previously been experienced separately, their preferences are initially unstable, but normally converge over the course of a session; silencing OFC activity impairs this update as well (Gardner *et al*., 2020). These results indicate that the OFC activity is crucial specifically at times when a behavioral strategy is being updated. They leave open the question of what computational role this activity plays. They are consistent with several possible roles, including identifying when the strategy should be updated, driving the update of that strategy directly, or driving decisions between recently-updated alternatives. Our analysis of the coarse effects of inactivation in different time windows replicates this result. Our analysis of the fine-grained effects of inactivation builds on it, using a cognitive model to jointly interpret neural recordings and trial-by-trial behavioral effects of inactivation. They indicate that the detailed effects of inactivation are attributable in particular to degradation of the outcome value signal (and not, for example, the reward signal, which is stronger and carried by a larger fraction of clusters). This suggests that, at least in our task, the OFC plays a particular computational role in supporting update: it provides outcome value estimates to a learning system, elsewhere in the brain, that updates the behavioral policy.

Our data are also broadly consistent with the idea that the OFC represents a “cognitive map” of task-relevant state information. A broad version of this account (Wilson et al., 2014) was based largely on lesion and inactivation data, and leaves open the question of whether this cognitive mapping information is used to drive choice directly, to guide learning, or for both. A narrower version of this cognitive mapping account was based also on fMRI data in human subjects (Chan, Niv and Norman, 2016; Schuck *et al*., 2016), and proposes that activity in the OFC specifically represents the subject’s beliefs about the current unobservable state of the environment. In our task, the most relevant states are fully observable (indicated by which noseports are currently lit). However it is possible to view the current reward probability contingencies (right/left outcome ports rewarded at 80%/20% vs. 20%/80%) as an unobservable task-relevant state variable that the animal might make inferences about. In our model, the “choice value difference” variable corresponds very closely to the current belief about this unobservable state. The narrow version of the cognitive mapping account might therefore predict that we would see this variable strongly represented in OFC throughout the entire trial. Our finding that it is represented only transiently and weakly puts our data in apparent tension with the narrower cognitive mapping account. A possible resolution to this tension lies in a difference in brain regions. The fMRI data (Chan, Niv and Norman, 2016; Schuck et al 2016) find their strongest representations of unobservable state in very medial regions of the orbital surface and in regions of the medial surface, likely belonging to the medial network (areas 10 and 11m; Price et al., 2007) rather than in the orbital network regions (areas 11l, 12/47, and 13) which are most plausibly homologous to the regions we investigated here (LO and AIv). A promising direction for future work would be to perform experiments analogous to ours in the regions of the rodent brain that are plausibly homologous to ventral parts of the medial network (VO and MO) as well as elsewhere in PFC

In summary, we find that rat behavior on a multi-step decision task dissociates the computational role of expected value information in choosing from its role in learning. We investigate the role of expected value representations in the OFC, and find a pattern of results indicating that these representations do not directly drive choice in this task, but instead support a learning process that updates choice mechanisms elsewhere in the brain. In addition to clarifying the function of the rodent OFC, these results provide insight into the overall architecture of value-based cognition. Specifically, they reveal that it is modular: that value-based learning and value-based choosing depend on separate neural representations. This lends credence to computational models which separate these functions (Joel, Niv and Ruppin, 2002; Song, Yang and Wang, 2017). It also motivates a search for the neural modules in the rat which play the remaining computational roles, including the role of directly driving choice.

## Author Contributions

KJM, MMB, and CDB conceived the project. KJM designed the experiments and analysis, with supervision from MMB and CDB. KJM carried out all experiments and analyzed the data. KJM and CDB wrote the manuscript, based on a first draft by KJM, with extensive comments from MMB.

## Acknowledgements

We thank Athena Akrami for assistance with array implant surgeries; Jovanna Teran, Klaus Osorio, Adrian Sirko, Samantha Stein, and Lillianne Teachen for assistance with animal training; and Peter Bibawi for help with histology. We thank Luca Mazzucato for identifying an error in an early version of our data processing pipeline. We thank Nathaniel Daw, Yael Niv, Geoffrey Schoenbaum, and Thomas Akam for helpful discussions, and Athena Akrami, Christine Constantinople, Nathaniel Daw, Cristina Domnisoru, Jeff Gauthier, Chuck Kopec, Olga Lositsky, Marcelo Mattar, Bas van Opheusden, Angela Radulescu, Ben Scott, Kim Stachenfeld, and Bob Wilson for helpful comments on the manuscript.

## Methods

### Subjects

All subjects were adult male Long-Evans rats (Taconic Biosciences; Hilltop Lab Animals), placed on a restricted water schedule to motivate them to work for water rewards. Rats were housed on a reverse 12-hour light cycle and trained during the dark phase of the cycle. Rats were pair housed during behavioral training and then single housed after being implanted with microwire arrays or optical fiber implants. Animal use procedures were approved by the Princeton University Institutional Animal Care and Use Committee and carried out in accordance with NIH standards. No explicit power analysis was used to determine the number of rats; instead we aimed for a sample size consistent with previous studies using similar tools.

### Two-Step Behavioral Task

Rats were trained on a two-step behavioral task, following a shaping procedure which has been previously described (Miller, Botvinick and Brody, 2017). Rats performed the task in custom behavioral chambers containing six “nose ports” arranged in two rows of three, each outfitted with a white LED for delivering visual stimuli, as well as an infrared LED and phototransistor for detecting rats’ nose entries into the port. The left and right ports in the bottom row also contained sipper tubes for delivering water rewards. The rat initiated each trial by entering the illuminated top center port, causing the two top side ports (“choice ports”) to illuminate. The rat then made his choice by entering one of these ports. Immediately upon entry into a choice port, two things happened: the bottom center port light illuminateed, and one of two possible sounds began to play, indicating which of the two bottom side ports (“outcome ports”) would eventually be illuminated. The rat then entered the bottom center port, which caused the appropriate outcome port to illuminate. Finally, the rat entered the outcome port which illuminated, and received either a water reward or an omission. Once the rat had consumed the reward, a trial-end sound played, and the top center port illuminated again to indicate that the next trial was ready. The selection of each choice port led to one of the outcome ports becoming available with 80% probability (common transition), and to the other becoming available with 20% probability (uncommon transition). These probabilities were counterbalanced across rats, but kept fixed for each rat for the entirety of his experience with the task. The probability that entry into each bottom side port would result in reward switched in blocks. In each block one port resulted in reward 80% of the time, and the other port resulted in reward 20% of the time. Block shifts happened unpredictably, with a minimum block length of 10 trials and a 2% probability of block change on each subsequent trial.

### Analysis of Behavioral Data: Planning Index & Model-Free Index

We quantify the effect of past trials and their outcomes on future decisions using a logistic regression analysis based on previous trials and their outcomes (Lau and Glimcher, 2005; Miller, Botvinick and Brody, 2017). We define vectors for each of the four possible trial outcomes: common-reward (CR), common-omission (CO), uncommon-reward (UR), and uncommon-omission (UO), each taking on a value of +1 for trials of their type where the rat selected the left choice port, a value of −1 for trials of their type where the rat selected the right choice port, and a value of 0 for trials of other types. We use the following regression model:

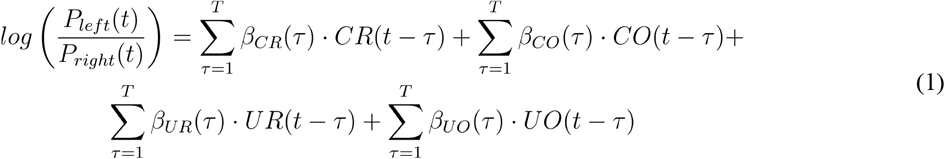

where *β*_*cr*_, *β*_*co*_, *β*_*ur*_, and *β*_*uo*_ are vectors of regression weights which quantify the tendency to repeat on the next trial a choice that was made τ trials ago and resulted in the outcome of their type, and *T* is a hyperparameter governing the number of past trials used by the model to predict upcoming choice, which was set to 3 for all analyses.

We expect model-free agents to repeat choices which lead to reward and switch away from those which lead to omissions (Daw *et al*., 2011), so we define a model-free index for a dataset as the sum of the appropriate weights from a regression model fit to that dataset:

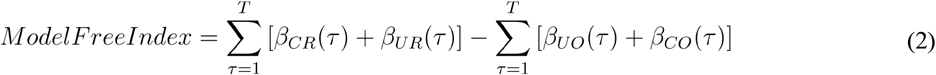

We expect that planning agents will show the opposite pattern after uncommon transition trials, since the uncommon transition from one choice is the common transition from the other choice. We define a planning index:

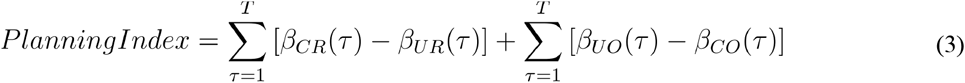

We also compute the main effect of past choices on future choice:

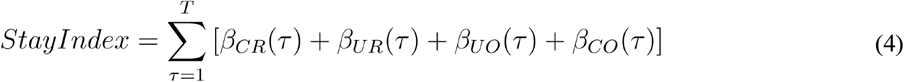

As well as an index quantifying the tendency to repeat choices the lead to common transitions and to switch away from those that lead to uncommon transitions:

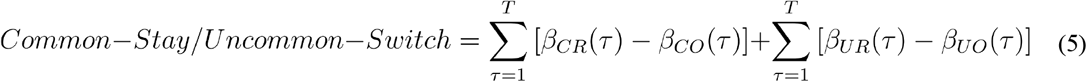

### Mixture-of-Agents Behavior Model

We model behavior and obtain trial-by-trial estimates of value signals using an agent-based computational model similar to one that we have previously shown to provide a good explanation of rat behavior on the two-step task (Miller, Botvinick and Brody, 2017). This model adopts the mixture-of-agents approach, in which each rat’s behavior is described as resulting from the influence of a weighted average of several different “agents” implementing different behavioral strategies to solve the task. On each trial, each agent *A* computes a value, *Q*_*A*_*(c)*, for each of the two available choices *c*, and the combined model makes a decision according to a weighted average of the various strategies’ values, *Q*_*total*_*(c)*:

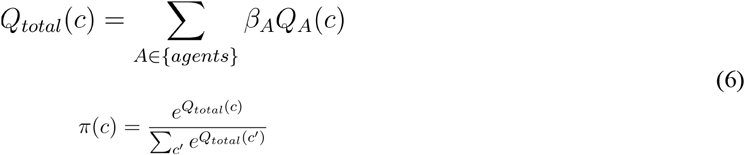

where the *β*’s are weighting parameters determining the influence of each agent, and π*(c)* is the probability that the mixture-of-agents will select choice *c* on that trial. The model which we have previously shown to provide the best explanation of rat’s behavior contains four such agents: model-based temporal difference learning, novelty preference, perseveration, and bias. The model used here is identical to the one in our previous paper (Miller et al. 2017), except that the perseverative agent is modified to allow it to consider many past trials rather than only the immediately previous trial (Miller et al. 2019).

#### Model-Based Temporal Difference Learning

Model-based temporal difference learning is a planning strategy, which maintains separate estimates of the probability with which each action (selecting the left or the right choice port) will lead to each outcome (the left or the right outcome port becoming available), *P(o*|*a)*, as well as the probability, *R(o)*, with which each outcome will lead to reward. This strategy assigns values to the actions by combining these probabilities to compute the expected probability with which selection of each action will ultimately lead to reward:

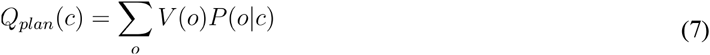

At the beginning of each session, the reward estimate *V(o)* is initialized to 0.5 for both outcomes, and the transition estimate *P(o*|*c)* is set to the true transition function for the rat being modeled (0.8 for common and 0.2 for uncommon transitions). After each trial, the reward estimate for both outcomes is updated according to

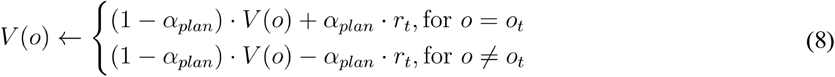

where *o*_*t*_ is the outcome that was observed on that trial, *r*_*t*_ is a binary variable indicating reward delivery, and *α* is a learning rate parameter constrained to lie between zero and one.

#### Novelty Preference

The novelty preference agent follows an *“*uncommon-stay/common switch” pattern, which tends to repeat choices when they lead to uncommon transitions on the previous trial, and to switch away from them when they lead to common transitions. Note that some rats have positive values of the *β*_*np*_ parameter weighting this agent (novelty preferring) while others have negative values (novelty averse; see **Fig 1e**):

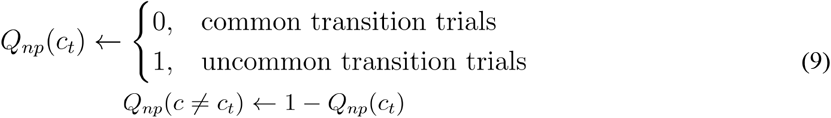

#### Perseveration

Perseveration is a pattern which tends to repeat choices that have been made in the recent past, regardless of whether they led to a common or an uncommon transition, and regardless of whether or not they led to reward.

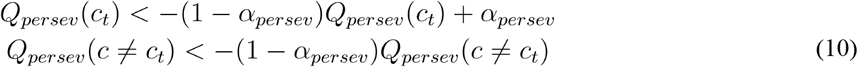

#### Bias

Bias is a pattern which tends to select the same choice port on every trial. Its value function is therefore static, with the extent and direction of the bias being governed by the magnitude and sign of this strategy’s weighting parameter *β*_*bias*_.

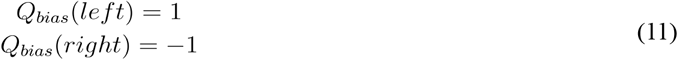

### Model Fitting

We implemented the model described above using the probabilistic programming language Stan (Carpenter *et al*., 2016; Stan Development Team, 2016), and performed maximum-a-posteriori fits using weakly informative priors on all parameters (Gelman *et al*., 2013) The prior over the weighting parameters *β* was normal with mean 0 and sd 0.5, and the prior over *α* was a beta distribution with *a*=*b*=3. For ease of comparison, we normalize the weighting parameters *β*_*plan*_, *β*_*np*_, and *β*_*persev*_, dividing each by the standard deviation of its agent’s associated values (*Q*_*plan*_, *Q*_*np*_, and *Q*_*persev*_) taken across trials. Since each weighting parameter affects behavior only by scaling the value output by its agent, this technique brings the weights into a common scale and facilitates interpretation of their relative magnitudes, analogous to the use of standardized coefficients in regression models.

### Surgery: Microwire Array Implants

Six rats were implanted with microwire arrays (Tucker-David Technologies) targeting OFC unilaterally. Arrays contained tungsten microwires 4.5mm long and 50 μm in diameter, cut at a 60° angle at the tips. Wires were arranged in four rows of eight, with spacing 250 μm within-row and 375 μm between rows, for a total of 32 wires in a 1.125 mm by 1.75 mm rectangle. Target coordinates for the implant with respect to bregma were 3.1-4.2mm anterior, 2.4-4.2mm lateral, and 5.2mm ventral (~4.2mm ventral to brain surface at the posterior-middle of the array).

In order to expose enough of the skull for a craniotomy in this location, the jaw muscle was carefully resected from the lateral skull ridge in the area near the target coordinates. Dimpling of the brain surface was minimized following procedures described in more detail elsewhere(Akrami *et al*., 2017). Briefly, a bolus of petroleum jelly (Puralube, Dechra Veterinary Products) was placed in the center of the craniotomy to protect it, while cyanoacrylate glue (Vetbond, 3M) was used to adhere the pia mater to the skull at the periphery. The petroleum jelly was then removed, and the microwire array inserted slowly into the brain. Rats recovered for a minimum of one week, with ad lib access to food and water, before returning to training.

### Electrophysiological Recordings

Once rats had recovered from surgery, recording sessions were performed in a behavioral chamber outfitted with a 32 channel recording system (Neuralynx). Spiking data was acquired using a bandpass filter between 600 and 6000 Hz and a spike detection threshold of 30 μV. Clusters were manually cut (Spikesort 3D, Neuralynx), and both single- and multi-units were considered. All manually-cut units were used for analysis. We observed that a small fraction of trials showed apparent artifacts in which some units appeared to have extremely high firing rates, which we suspect was due to motion of the implant or tether. We therefore excluded units from analysis on trials where the Median Absolute Deviation (Leys *et al*., 2013) of their spike count exceeded a conservative threshold of three.

### Analysis of Electrophysiology Data

To determine the extent to which different variables were encoded in the neural signal, we fit a series of regression models to our spiking data. Models were fit to the spike counts emitted by each unit in 200 ms time bins taken relative to the four noseport entry events that made up each trial. There were ten total regressors, defined relative to a pair of adjacent trials. Seven of them were binary (coded as +-1), and related to observable task variables: the choice port selected on the earlier trial(left or right), the outcome port visited (left or right), the reward received (reward or omission), the interaction between choice port and outcome port (common or uncommon transition), the interaction between choice port and reward, the interaction between outcome port and reward, and the choice port selected on the later trial. Three additional regressors were continuous and related to subjective reward expectation: the expected value of the outcome port visited (*V*) for the first trial, the value difference between the choice ports (*Q(left)* - *Q(right)*), and the value of the choice port selected (*Q(chosen)*) on the subsequent trial. These last three regressors were obtained using the agent-based computational model described above, with parameters fit separately to each rat’s behavioral data. Regressors were z-scored to facilitate comparison of fit regression weights. Models were fit using the Matlab function `glmnet` (Qian *et al*., 2013) using a Poisson noise model. We fit each model both with no regularization and with L1 regularization (*α* = 1, λ = 0, 10^−10^, 10^−9^, …, 10^−1^), using the model with the weakest regularization that still allowed all weights to be identifiable.

In our task, many of these regressors were correlated with one another (**Fig3-S2**), so we quantify encoding using the coefficient of partial determination (CPD; also known as partial r-squared) associated with each (Cai, Kim and Lee, 2011; Kennerley, Behrens and Wallis, 2011). This measure quantifies the fraction of variance explained by each regressor, once the variance explained by all other regressors has been taken account of:

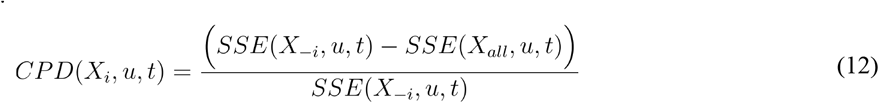

where *u* refers to a particular unit, *t* refers to a particular time bin, and SSE(*X*_*all*_) refers to the sum-squared-error of a regression model considering all eight regressors described above, and SSE(X_-i_) refers to the sum-squared-error of a model considering the seven regressors other than *X*_*i*_. We compute total CPD for each unit by summing the SSE associated with the regression models for that unit for all time bins:

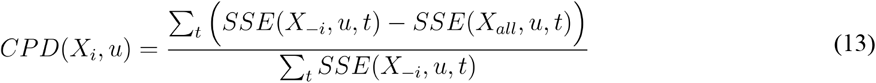

We report this measure for individual example units showing all time bins (Fig 2, bottom). We report the CPD for each unit for particular port entry events (Fig 3a), taking the sum over the five bins making up a 1s time window centered on a particular port entry event (top neutral center port, choice port, bottom neutral center port, or outcome port). We report the “population CPD” (Fig 3b, Fig 3c) by aggregating over all units for a particular time bin:

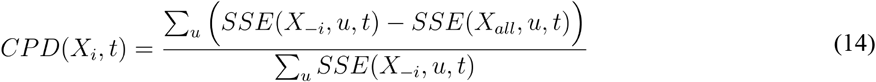

For each of these measures, we assess significance by comparing the CPD computed on the true dataset to a distribution of CPDs computed on surrogate datasets constructed by circularly permuting the trial labels within each session. We use permuted, rather than shuffled, labels in order to preserve trial-by-trial correlational structure. If the true CPD is larger than most of the CPDs from these surrogate datasets, we can reject the null hypothesis that the CPD is driven by correlational structure alone. We compute a permutation p-value by finding the percentile of the true CPD within the distribution of CPDs in surrogate datasets. In the main-text figure (Figure 3b, Figure 3c) we additionally subtract the mean of the CPDs from the surrogate datasets, in order to give a measure that can be fairly compared to zero.

### Surgery: Optical Fiber Implant and Virus Injection

Rats were implanted with sharpened fiber optics and received virus injections following procedures similar to those described previously (Hanks *et al*., 2015; Kopec *et al*., 2015; Akrami *et al*., 2017), and documented in detail on the Brody lab website (http://brodywiki.princeton.edu/wiki/index.php/Etching_Fiber_Optics). A 50/125 μm LC-LC duplex fiber cable (Fiber Cables) was dissected to produce four blunt fiber segments with LC connectors. These segments were then sharpened by immersing them in hydroflouric acid and slowly retracting them using a custom-built motorized jig attached to a micromanipulator (Narashige International) holding the fiber. Each rat was implanted with two sharpened fibers, in order to target OFC bilaterally. Target coordinates with respect to bregma were 3.5mm anterior, 2.5mm lateral, 5mm ventral. Fibers were angled 10 degrees laterally, to make space for female-female LC connectors which were attached to each and included as part of the implant.

Four rats were implanted with sharpened optical fibers only, but received no injection of virus. These rats served as uninfected controls.

Nine additional rats received both fiber implants as well as injections of a virus (AAV5-CaMKII*α*-eNpHR3.0-eYFP; UNC Vector Core) into the OFC to drive expression of the light-activated inhibitory opsin eNpHR3.0. Virus was loaded into a glass micropipette mounted into a Nanoject III (Drummond Scientific), which was used for injections. Injections involved five tracks arranged in a plus-shape, with spacing 500μm. The center track was located 3.5mm anterior and 2.5mm lateral to bregma, and all tracks extended from 4.3 to 5.7mm ventral to bregma. In each track, 15 injections of 23 nL were made at 100μm intervals, pausing for ten seconds between injections, and for one minute at the bottom of each track. In total 1.7 μl of virus were delivered to each hemisphere over a period of about 20 minutes.

Rats recovered for a minimum of one week, with ad lib access to food and water, before returning to training. Rats with virus injections returned to training, but did not begin inactivation experiments until a minimum of six weeks had passed, to allow for virus expression.

### Optogenetic Perturbation Experiments

During inactivation experiments, rats performed the task in a behavioral chamber outfitted with a dual fiber optic patch cable connected to a splitter and a single-fiber commutator (Princetel) mounted in the ceiling. This fiber was coupled to a 200 mW 532 nm laser (OEM Laser Systems) under the control of a mechanical shutter (ThorLabs) by way of a fiber port (ThorLabs). The laser power was tuned such that each of the two fibers entering the implant received between 25 and 30 mW of light when the shutter was open. Each rat received several sessions in which the shutter remained closed, in order to acclimate to performing the task while tethered. Once the rat showed behavioral performance while tethered that was similar to his performance before the implant surgery, inactivation sessions began. During these sessions, the laser shutter was opened (causing light to flow into the implant, activating the eNpHR3.0 and silence neural activity) on 7% of trials each in one of three time periods. “Outcome period” inactivation began when the rat entered the bottom center port at the end of the trial, and ended either when the rat had left the port and remained out for a minimum of 500 ms, or after 2.5 s. “Choice period” inactivation began at the end of the outcome period and lasted until the rat entered the choice port on the following trial. “Both period” inactivation encompassed both the outcome period and the choice period. The total duration of the inactivation therefore depended in part on the movement times of the rat, and was somewhat variable from trials to trial (**Fig 5-S2**). If a scheduled inactivation would last more than 15 s, inactivation was terminated, and that trial was excluded from analysis. Due to constraints of the bControl software, inactivation was only performed on even-numbered trials.

### Analysis of Optogenetic Effects on Behavior

We quantify the effects of optogenetic inhibition on behavior by computing separately the planning index for trials following inactivation of each type (outcome period, choice period, both periods) and for control trials. Specifically, we fit the trial history regression model of Equation 1 with a separate set of weights for trials following inactivation of each type:

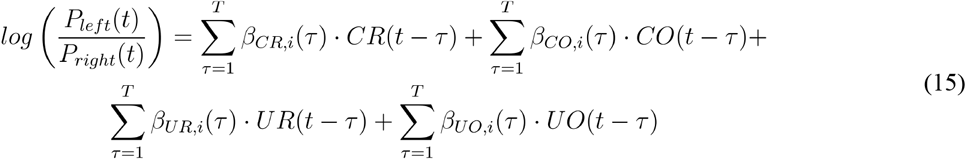

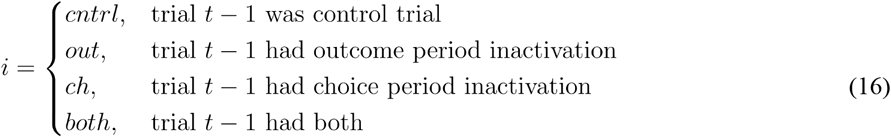

We used maximum a posteriori fitting in which the priors were Normal(0,1) for weights corresponding to control trials, and Normal(*β*_*X, cntrl*_, 1) for weights corresponding to inactivation trials, where *β*_*X, cntrl*_ is the corresponding control trial weight – e.g. the prior for *β*_*CR, out*_(1) is Normal(*β*_*CR, cntrl*_(1), 1). This prior embodies the belief that the effect of inactivation on behavior is likely to be small, and that the direction of any effect is equally likely to be positive or negative. This ensures that our priors cannot induce any spurious differences between control and inactivation conditions into the parameter estimates. We then compute a planning index separately for the weights of each type, modifying equation 3:

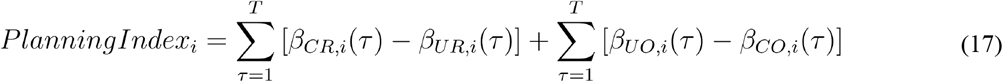

We compute the relative change in planning index for each inactivation condition: (*PlanningIndex*_*i*_ - *PlanningIndex*_*cntrl*_) / *PlanningIndex*_*cntrl*_, and report three types of significance tests on this quantity. First, we test for each inactivation condition the hypothesis that there was a significant change in the planning index, reporting the results of a one-sample t-test over rats. Next, we test the hypothesis that different inactivation conditions had effects of different sizes on the planning index, reporting a paired t-test over rats. Finally, we test the hypothesis for each condition that inactivation had a different effect than sham inactivation (conducted in rats which had not received virus injections to deliver eNpHR3.0), reporting a two-sample t-test.

To test the hypothesis that inactivation specifically impairs the effect of distant past outcomes on upcoming choice, we break down the planning index for each condition by the index of the weights the contribute to it:

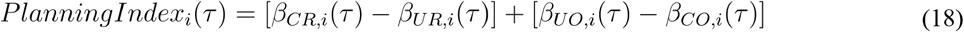

We report these trial-lagged planning indices for each inactivation condition, and assess the significance of the difference between inactivation and control conditions at each lag using a paired t-test.

### Synthetic Datasets

To generate synthetic datasets for comparison to optogenetic inactivation data, we generalized the behavioral model to separate the contributions of representations of expected value and of immediate reward. In particular, we replaced the learning equation within the model-based RL agent (Equation 7) with the following:

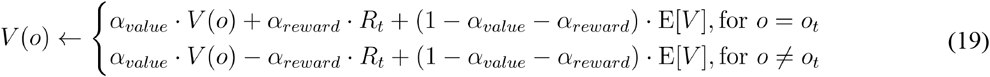

where *α*_*value*_ and *α*_*reward*_ are separate learning rate parameters, constrained to be nonnegative and to have a sum no larger than one, and E[*V*] represents the expected reward of a random-choice policy on the task, which in the case of our task is equal to 0.5.

To generate synthetic datasets in which silencing the OFC impairs choice value representations, outcome value representations, or reward representations, we decrease the parameter *β*_*plan*_, *α*_*value*_, or *α*_*reward*_, respectively. Specifically, we first fit the model to the dataset for each rat in the optogenetics experiment (n=9) as above (i.e. using equation 7 as the learning rule) to obtain maximum a posteriori parameters. We translated these parameters to the optogenetics (equation 18) version of the model by setting *α*_*value*_ equal to the fit parameter *α* and *α*_*reward*_ equal to 1 - *α*. We then generated four synthetic datasets for each rat. For the control dataset, the fit parameters were used on trials of all types, regardless of whether inhibition of OFC was scheduled on that trial. For the “impaired outcome values” dataset, *α*_*value*_ was decreased specifically for trials with inhibition scheduled during the outcome period or both periods, but not on trials with inhibition during the choice period or on control trials. For the “impaired reward processing” dataset, *α*_*reward*_ was decreased on these trials instead. For the “impaired decision-making” dataset, *β*_*plan*_ was decreased specifically on trials following inhibition. In all cases, the parameter to be decreased was multiplied by 0.3, and synthetic datasets consisted of 100,000 total trials per rat.

### Histological Verification of Targeting

We verified that surgical implants were successfully placed in the OFC using standard histological techniques. At experimental endpoint, rats with electrode arrays were anesthetized, and microlesions were made at the site of each electrode tip by passing current through the electrodes. Rats were then perfused transcardially with saline followed by formalin. Brains were sliced using a vibratome and imaged using an epifluorescent microscope. Recording sites were identified using these microlesions and the scars created by the electrodes in passing, as well dimples in the surface of the brain. Locations of optical fibers were identified using the scars created by their passage. Location of virus expression was identified by imaging the YFP conjugated to the eNpHR3.0 molecule. **Figure 5-S1**

### Data and Code Availability

Data collected for the purpose of this paper will be posted on Figshare upon acceptance. Software used to analyze the data will be made available as a Github release. Software used for training rats and design files for constructing behavioral rigs are available on the Brody lab website.

---

**Figure 2 - Supplement 1:**
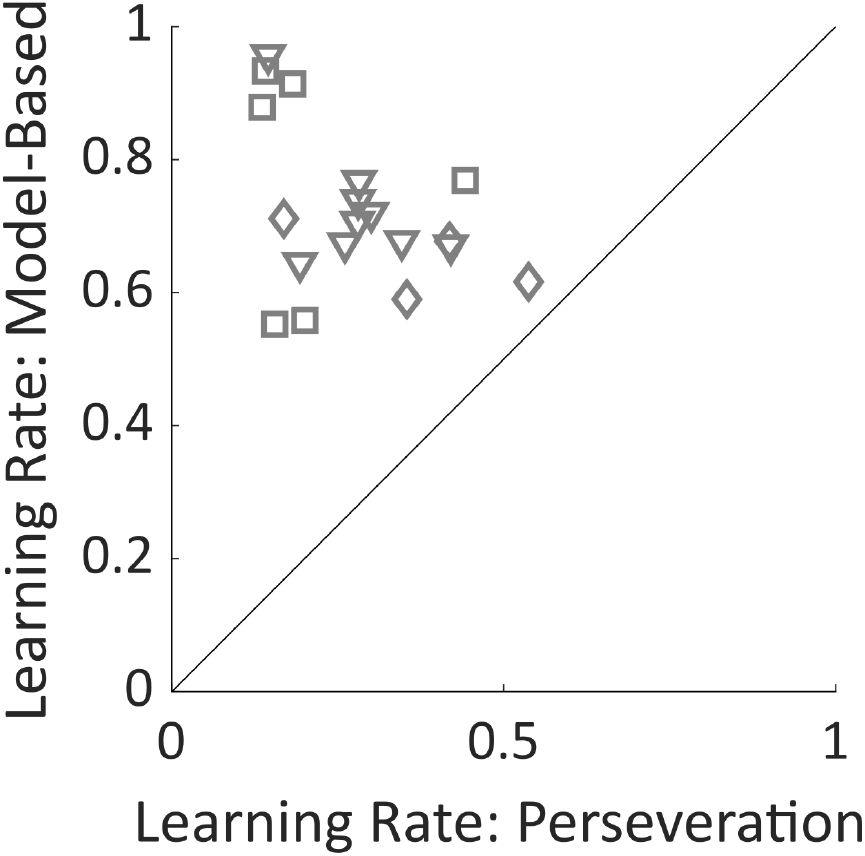
Learning Rate Parameters. Fit learning rate parameters from the mixture-of-agents model for the model-based planning agent and for the perseverative agent. Learning rates for perseveration are smaller for all rats, indicating that this agent takes into account a larger number of recent trials than the model-based agent does. Shapes of symbols indicate rats participating in the electrophysiology (squares), optogenetics (triangles) or sham optogenetics (diamonds) experiments.

**Figure 2 - Supplement 2:**
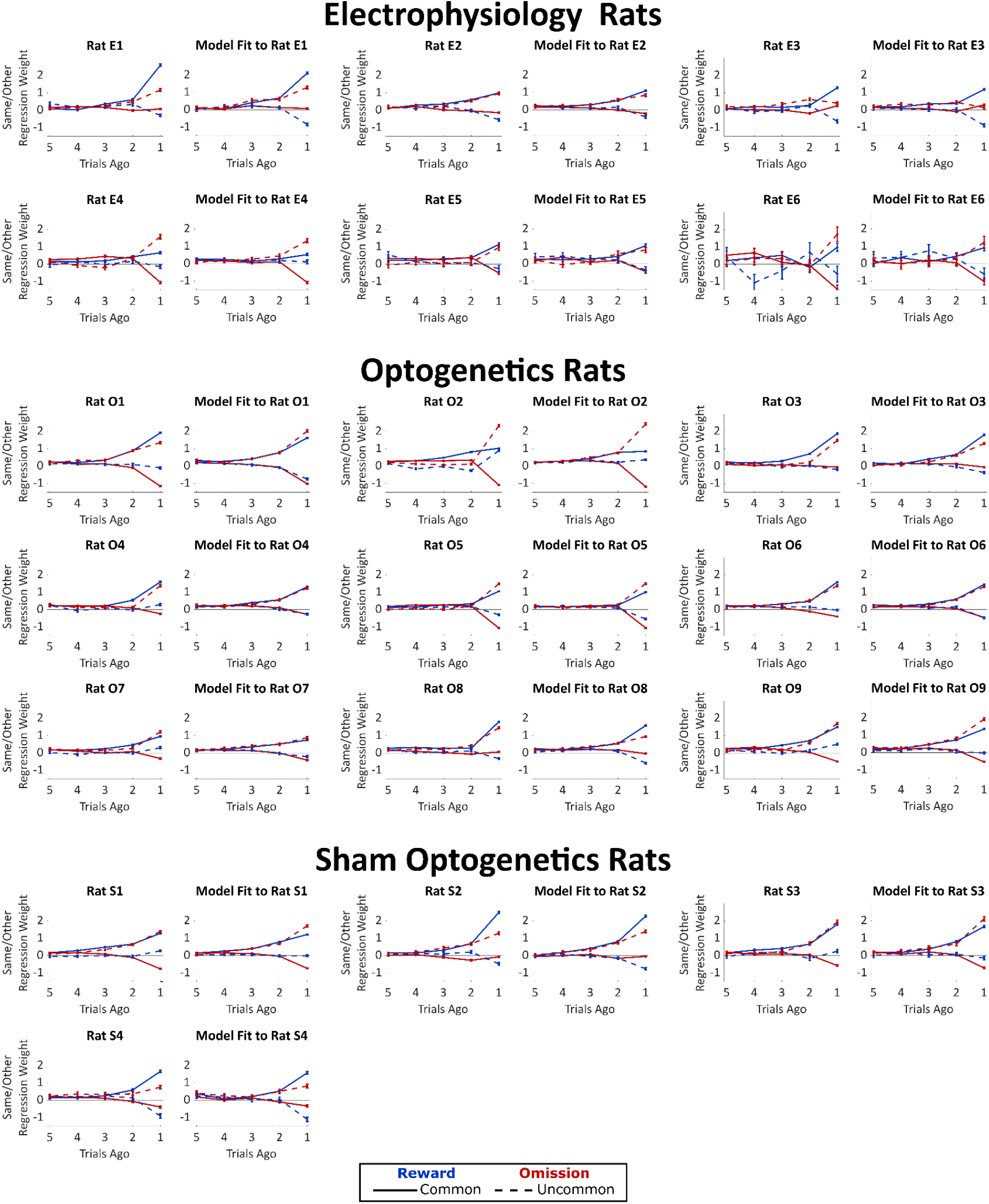
Weights of trial-history regression model, fit both to rats’ behavioral data and to synthetic datasets generated by our mixture-of-agents model fit separately to each rat. The planning strategy is characterized by a greater tendency to repeat choices that lead to rewards following a common transition than following an uncommon transition (solid blue line above dotted blue line), as well as a greater tendency to repeat choices that lead to omissions following an uncommon transition than following a common transition (dotted red line above solid red line). All nineteen rats show evidence of such a strategy.

**Figure 3 - Supplement 1:**
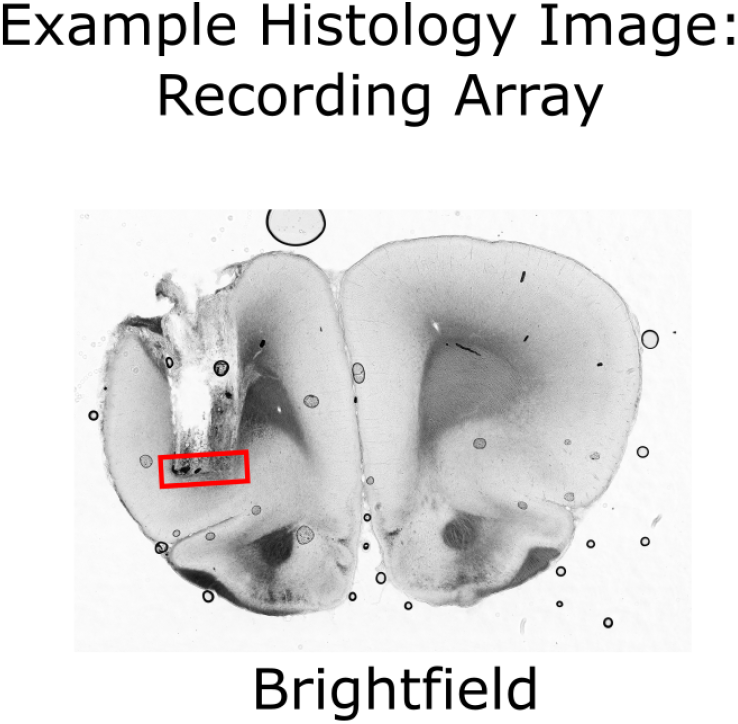
Histological verification of implant locations in OFC. Brightfield image of a coronal section from a rat implanted with an electrode array targeting OFC. The locations of the electrode tips are visible, as is damage done when the array was removed post-mortem. Red box indicates the estimated location of electrode tips.

**Figure 3 - Supplement 2:**
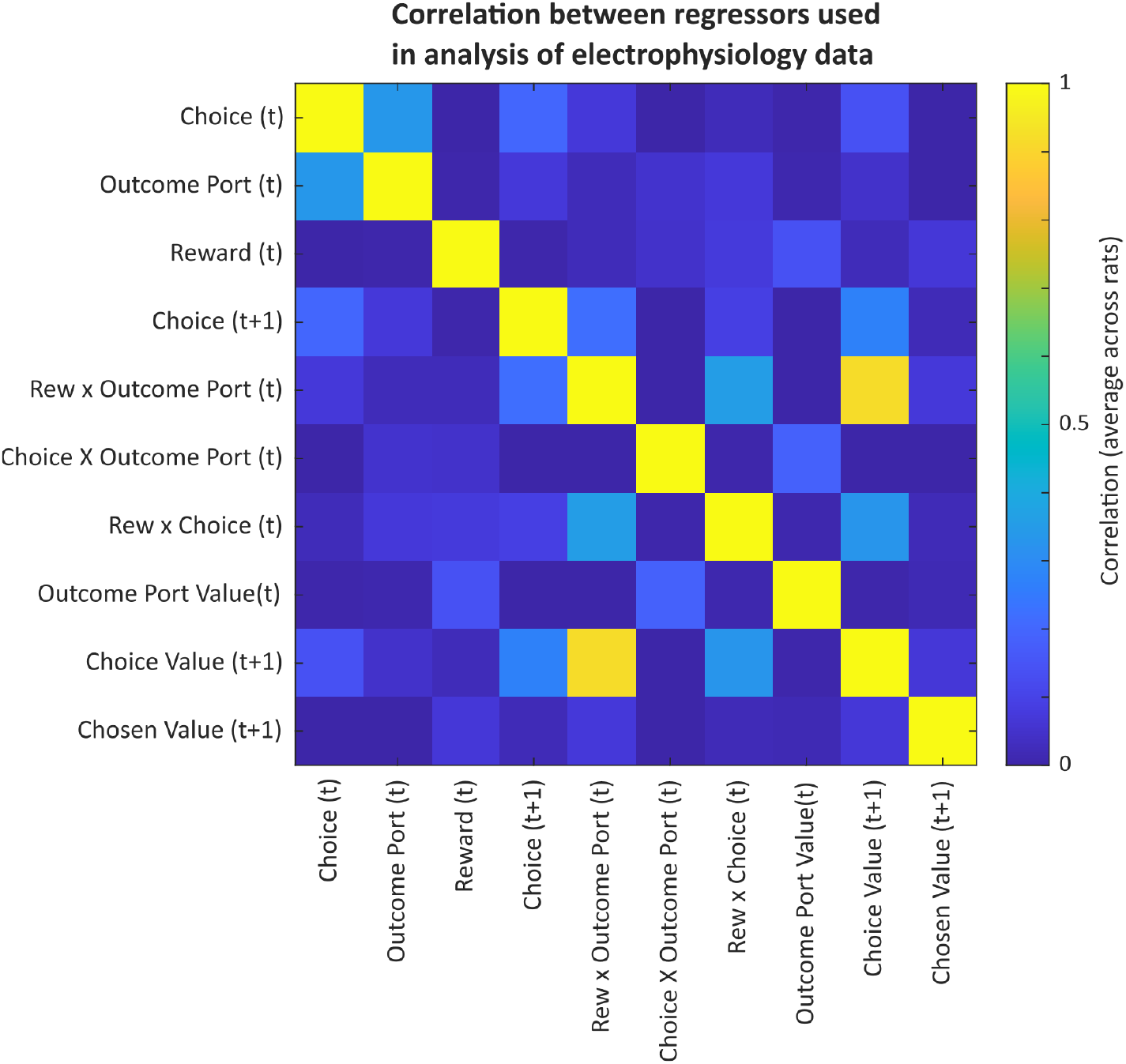
Correlations among predictors in the model used for analysis of electrophysiology data. Correlations among regressors motivate the use of a coefficient of partial determination analysis to quantify the unique contribution of each predictor to explaining neural activity.

**Figure 4 - Supplement 1:**
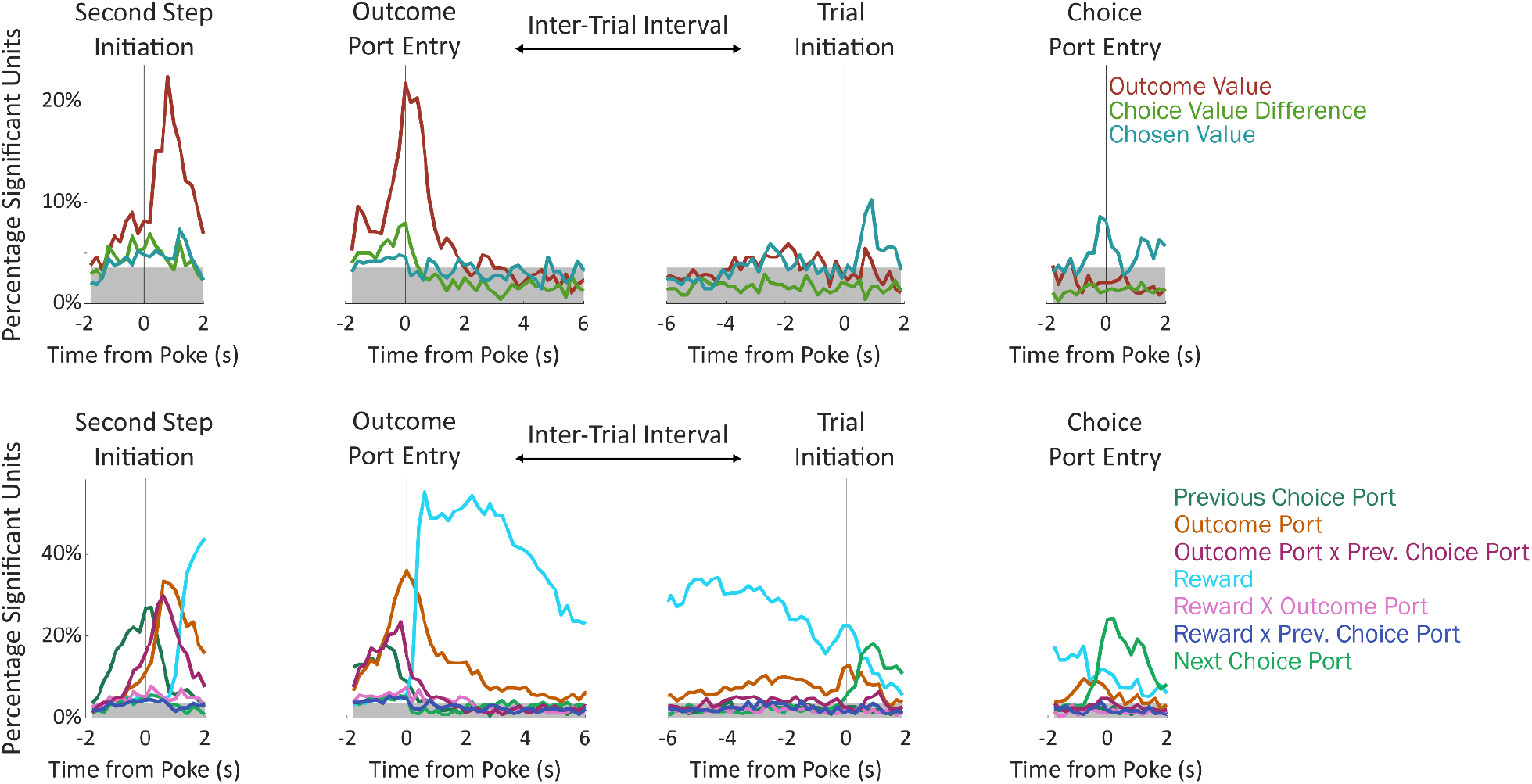
Fraction of units significantly encoding each regressor in each time bin. Units were deemed significant if they earned a coefficient of partial determination larger than that of 99% of permuted datasets for that regressor in that time bin. Gray shading indicates a threshold for population-level significance (17/477 units; equivalent to p=2×10^−6^ by a Bernoulli test uncorrected; p<0.01 after Bonferroni correction)

**Figure 4 - Supplement 2:**
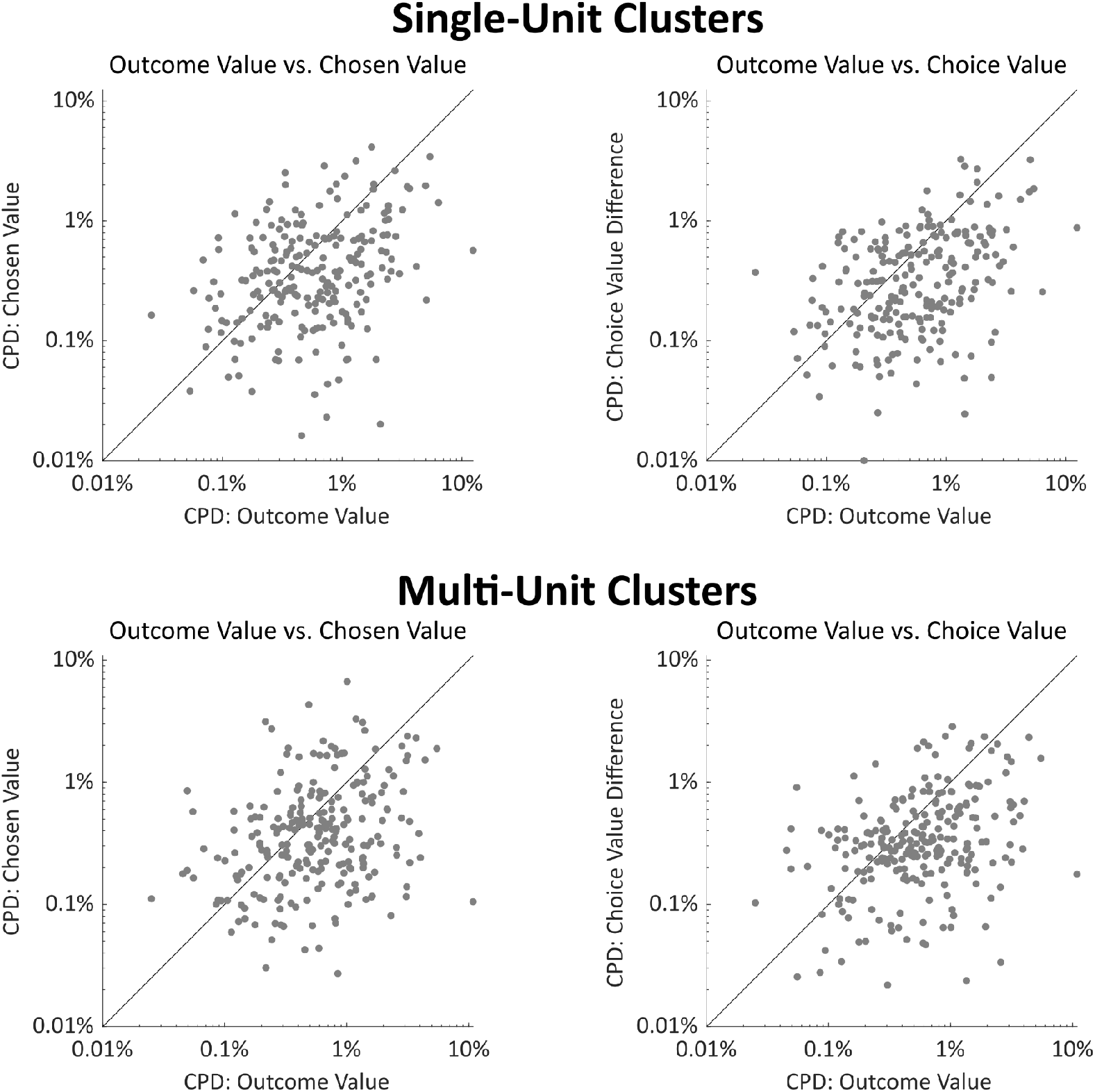
**above.** Right: Scatterplot showing CPD for each single unit (n=251) for the outcome-value regressor against CPD for the choice-value-difference regressor, both computed in a one-second window centered on entry into the outcome port. Right: Scatterplot showing CPD for outcome-value, computed at outcome port entry, against CPD for the chosen-value regressor, computed at choice port entry. **below**. As above but for multi-unit clusters (n=226). In all panels, CPD for outcome value is greater than for choice-related value information (all p<10^−7^, signrank test. See also Fig 4a).

**Figure 4 - Supplement 3:**
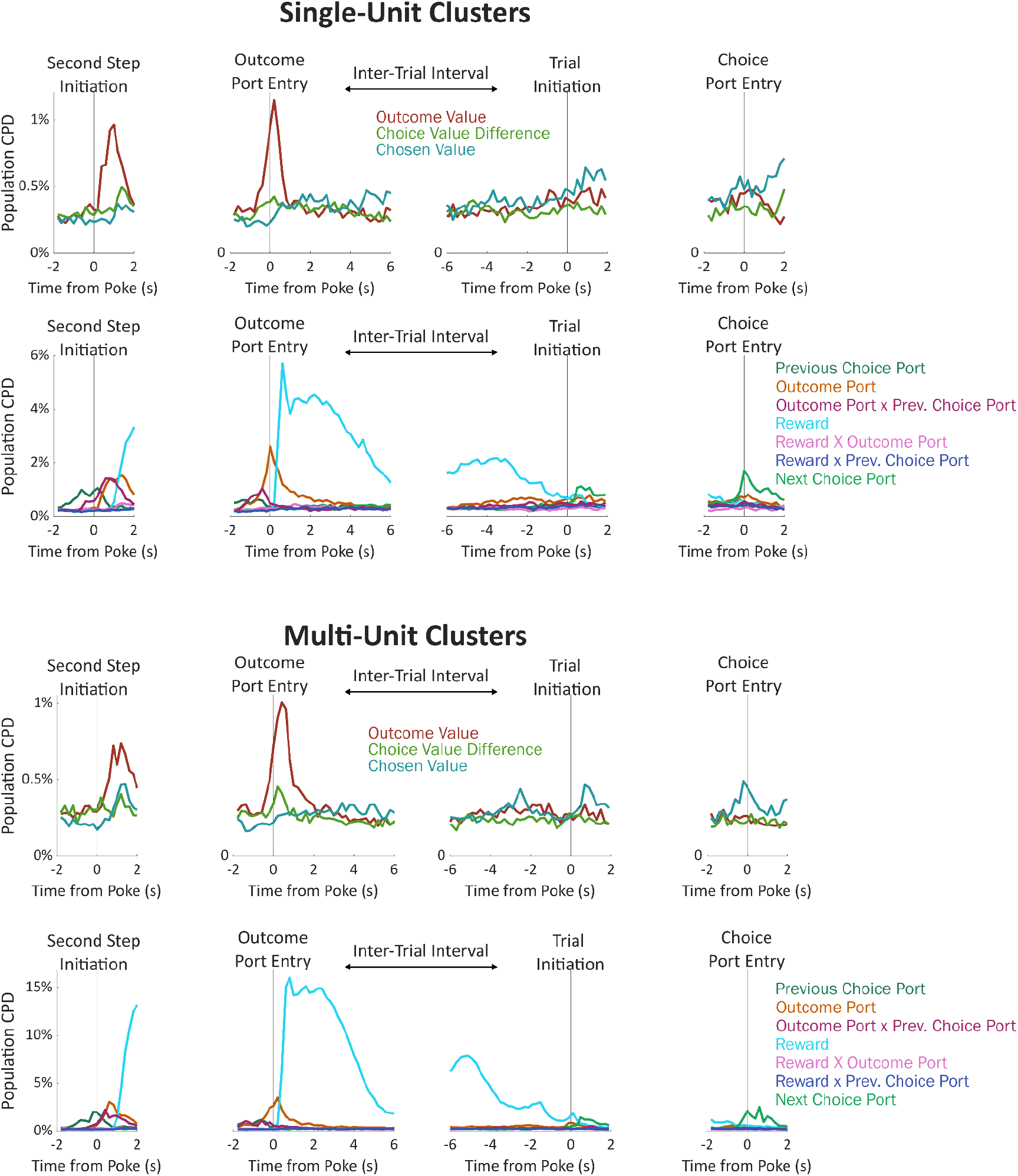
Timecourse of population CPD for the regressors in our model, considering only single-unit clusters (above) or considering only multi-unit clusters (below). See also Fig. 2a,c.

**Figure 4 - Supplement 4:**
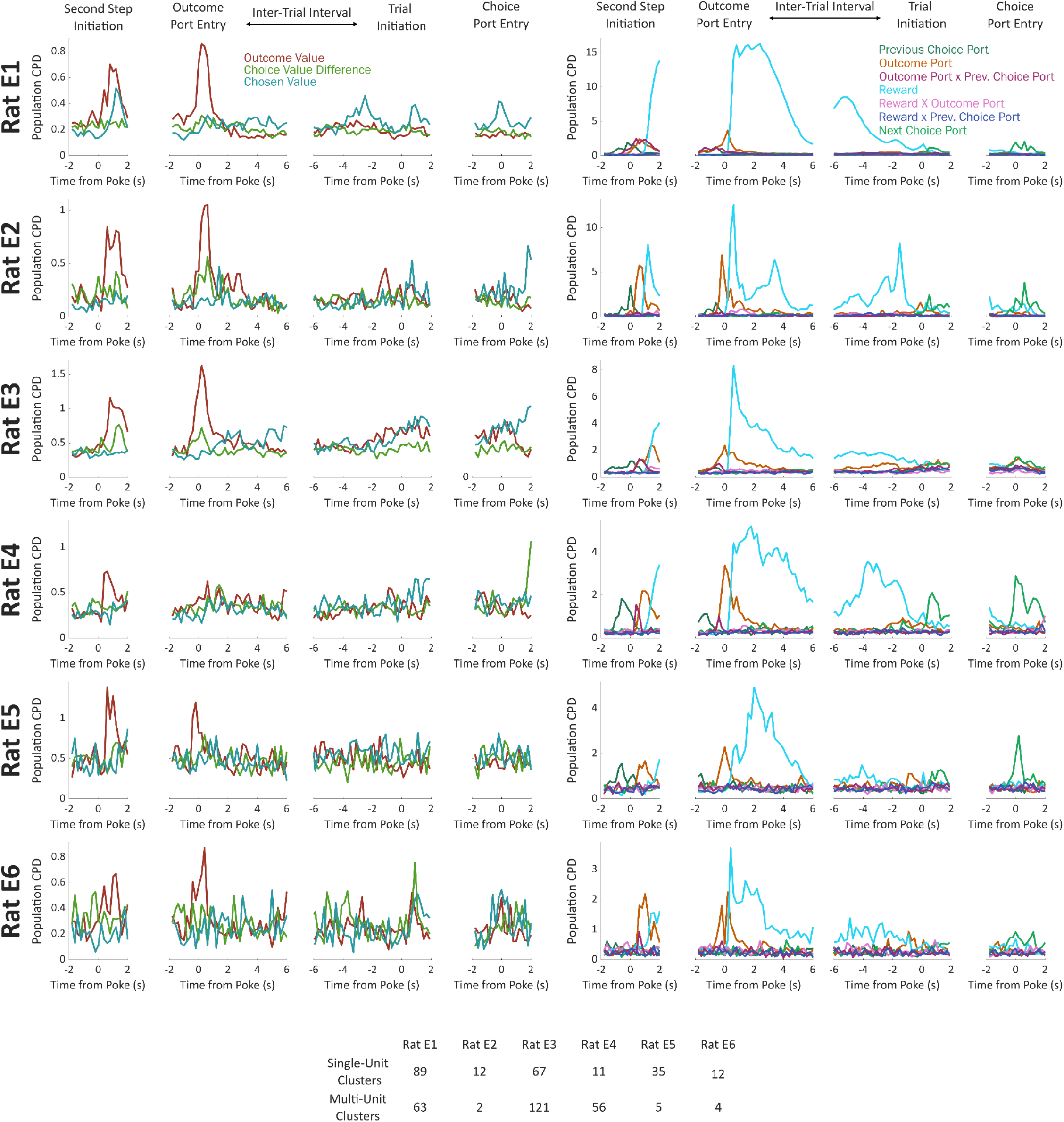
Analysis considering each rat individually. Above: Population CPD computed separately for units from each rat in the dataset. Below: Number of single-unit and number of multi-unit clusters recorded in each rat.

**Figure 4 - Supplement 5:**
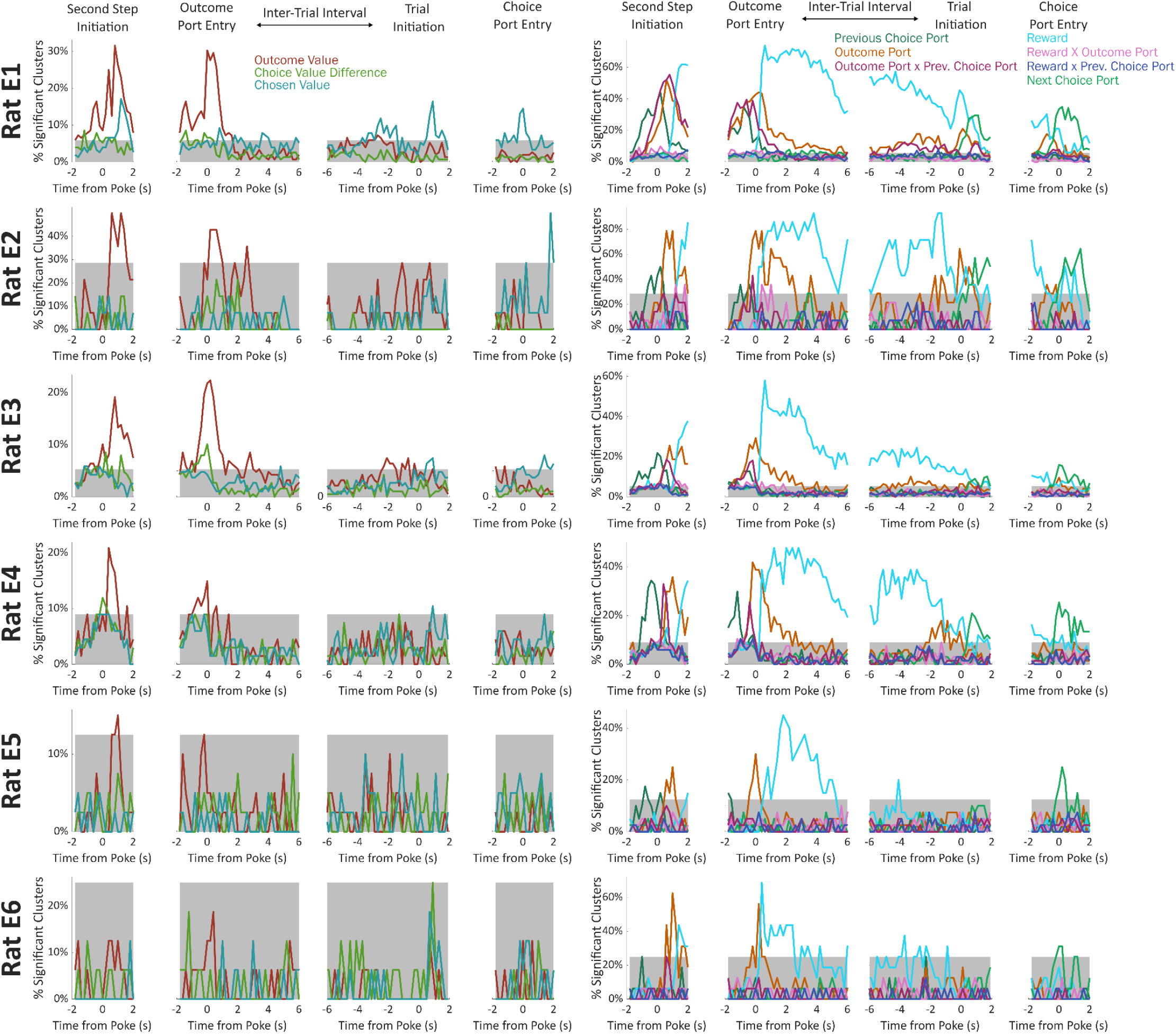
Fraction of significant units, considering each rat individually. Units were deemed significant if they earned a coefficient of partial determination larger than that of 99% of permuted datasets for that regressor in that time bin. Gray shading indicates a threshold for rat-level significance (p=2×10^−6^ by a Bernoulli test uncorrected; p<0.01 after Bonferroni correction)

**Figure 4 - Supplement 6:**
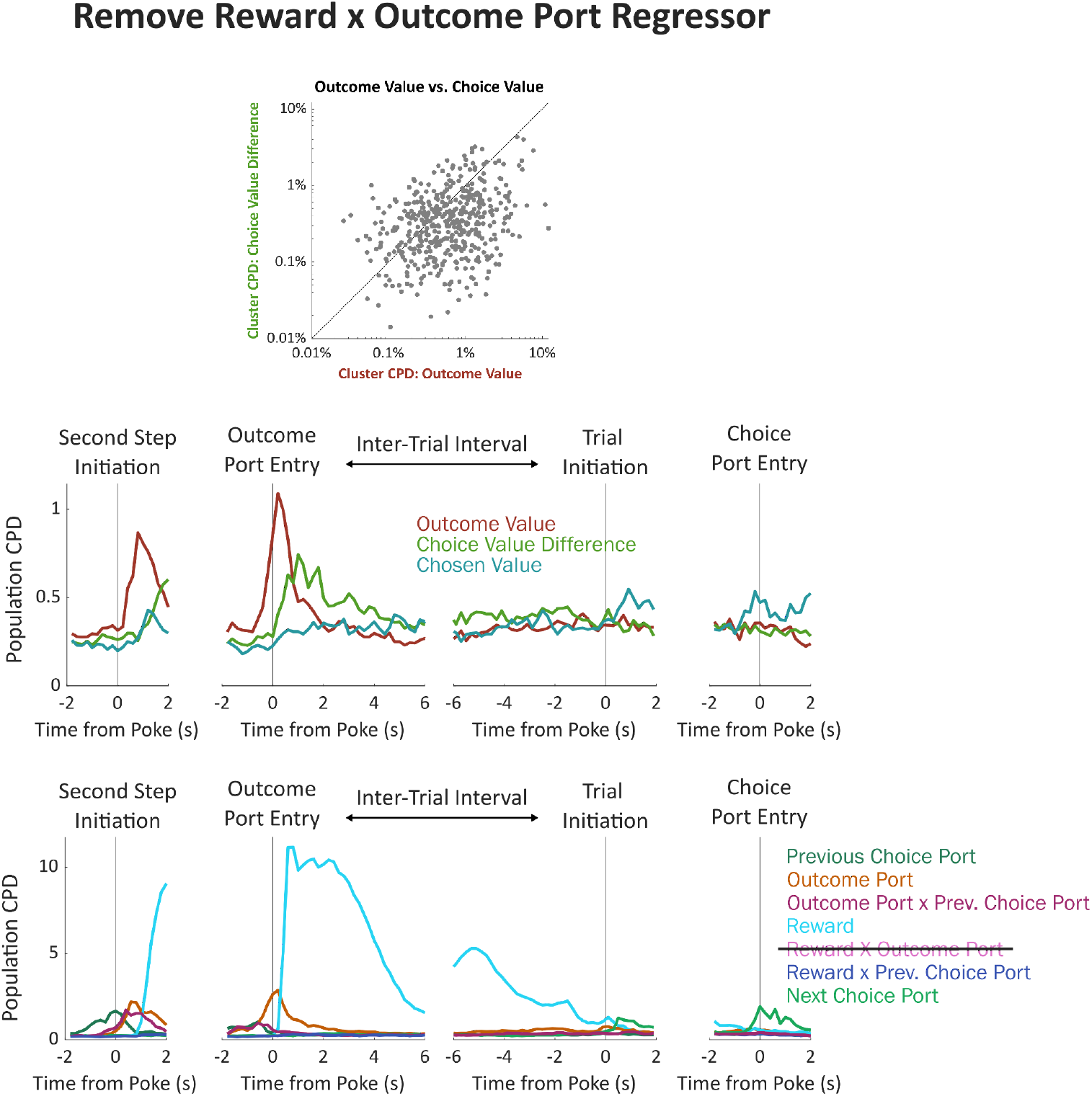
Analysis removing the outcome-by-reward interaction regressor. The coefficient of partial determination earned by each regressor is sensitive to our choice of which other regressors to include in the model. The strength of this sensitivity depends on the correlation between the regressors (adding an orthogonal regressor will not affect CPD; adding a perfectly correlated regressor will reduce CPD to zero). The “choice value difference” regressor in our model is highly correlated with one of our other regressors “outcome port by reward interaction”. This is because, in our model, outcome port and reward information are used to update outcome port values, which in turn update choice port values: a reward at the left outcome port or an omission at the right outcome port will increase the relative value of one choice port; a reward at the right outcome port or an omission at the left outcome port will increase the relative value of the other. This raises the possibility that our finding that the choice value difference regressor earns a relatively small CPD is an artifact of this correlation. To check this, we re-ran our regression analysis without the “outcome port by reward interaction” regressor. Plots of individual clusters (**a**) and of the population timecourse (**b**) show that choice value difference still earns ambler CPDs than outcome value.

**Figure 4 - Supplement 7:**
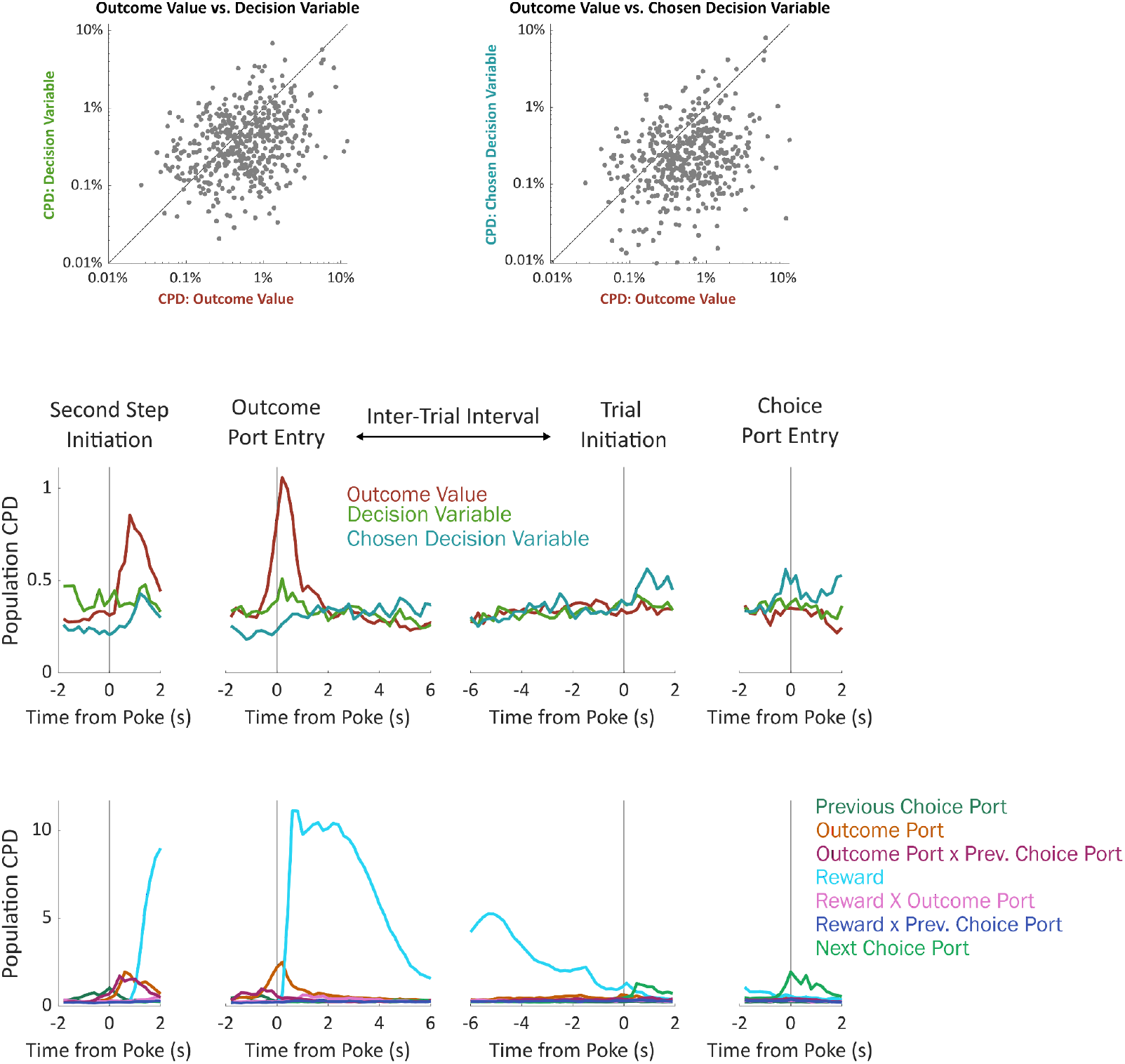
Analysis using alternative choice-related value regressors. To compute choice-related value regressors, we used the expected values of the model-based planning component of our cognitive model (*Q*, see Figure 2a). However the decision variable used by our model is not this expected value in isolation, but instead a weighted sum including expected value as well as variables related to perseveration and novelty preference (*P* and *N*, Figure 2a). Here, we compute analogs of choice value difference and of chosen value in terms of this decision variable instead of in terms of Q.

**Figure 4 - Supplement 8:**
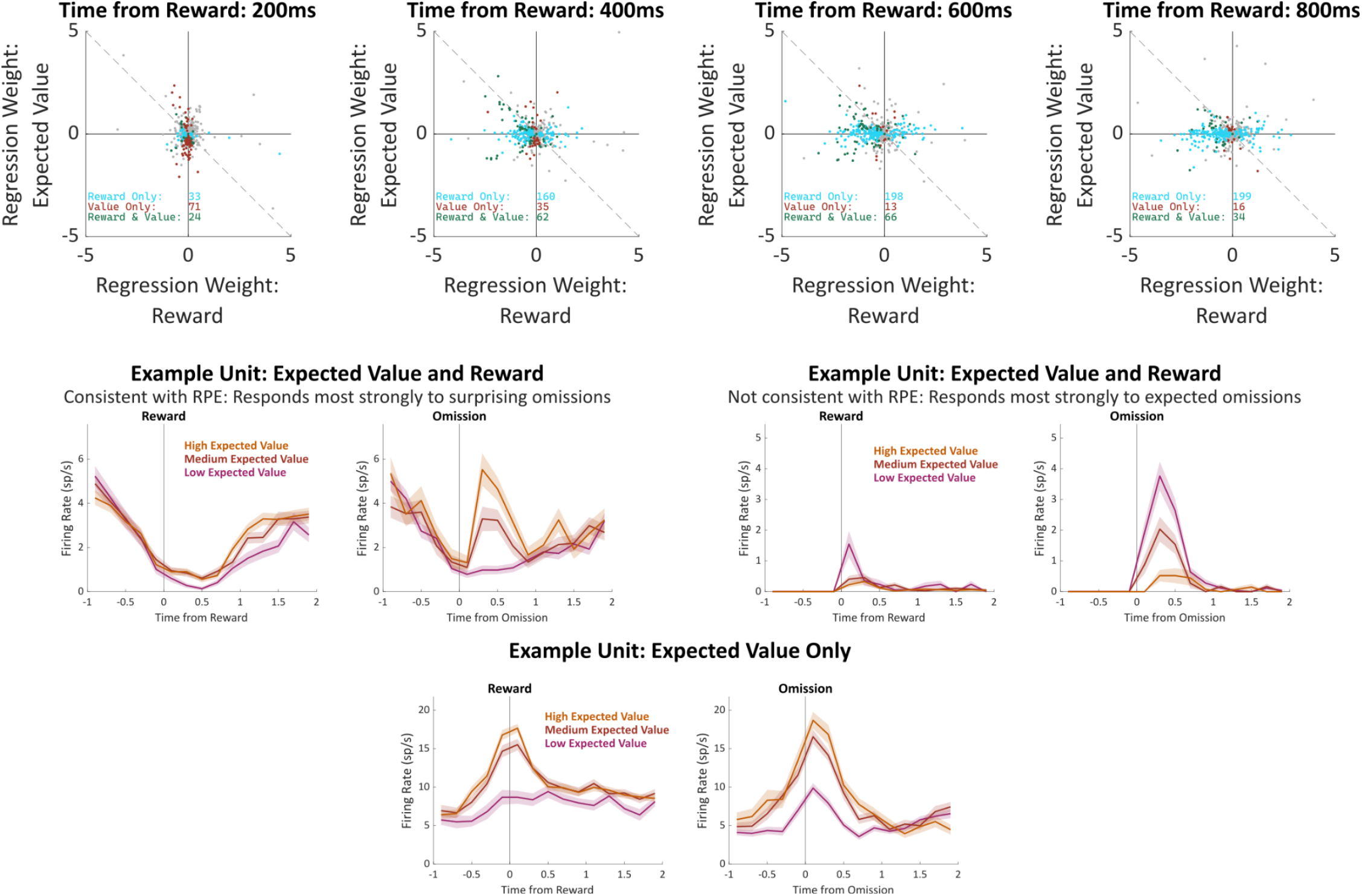
Correlates of reward prediction error. It has been suggested that OFC causes value-guided learning in other brain regions via signaling of reward prediction errors (RPE; Banerjee *et al*., 2020). RPE is defined as the difference between the reward that was actually experienced and the reward that was expected. **above**: We tested whether our OFC units tended to carry such a signal by examining the fit weights from our regression model in the time bins immediately following reward, following the method of Sul et al., (2010). A unit signaling RPE in our task is expected to have equal and opposite regression weights for reward and for outcome port expected value. Considering all units, we find that there is a weak but significant tendency for these weights to have opposite sign in two of the time bins (400ms: 257/477, 54%, p=0.04; 600ms: 262/477, 55%, p=0.01). Considering only units which correlate significantly with both reward and expected value, there was a similar trend (400ms: 37/62, 60%, p=0.05; 600ms: 39/66, 59%, p=0.05). These results are consistent with the idea that OFC contains units which correlate with RPE, with the caveats that it also contains a substantial number of units which correlate with its opposite (reward expected *plus* reward received), as well as units which correlate with expected reward without correlating with reward itself at all. **Below**: Example units illustrating these three patterns.

**Figure 5 - Supplement 1:**
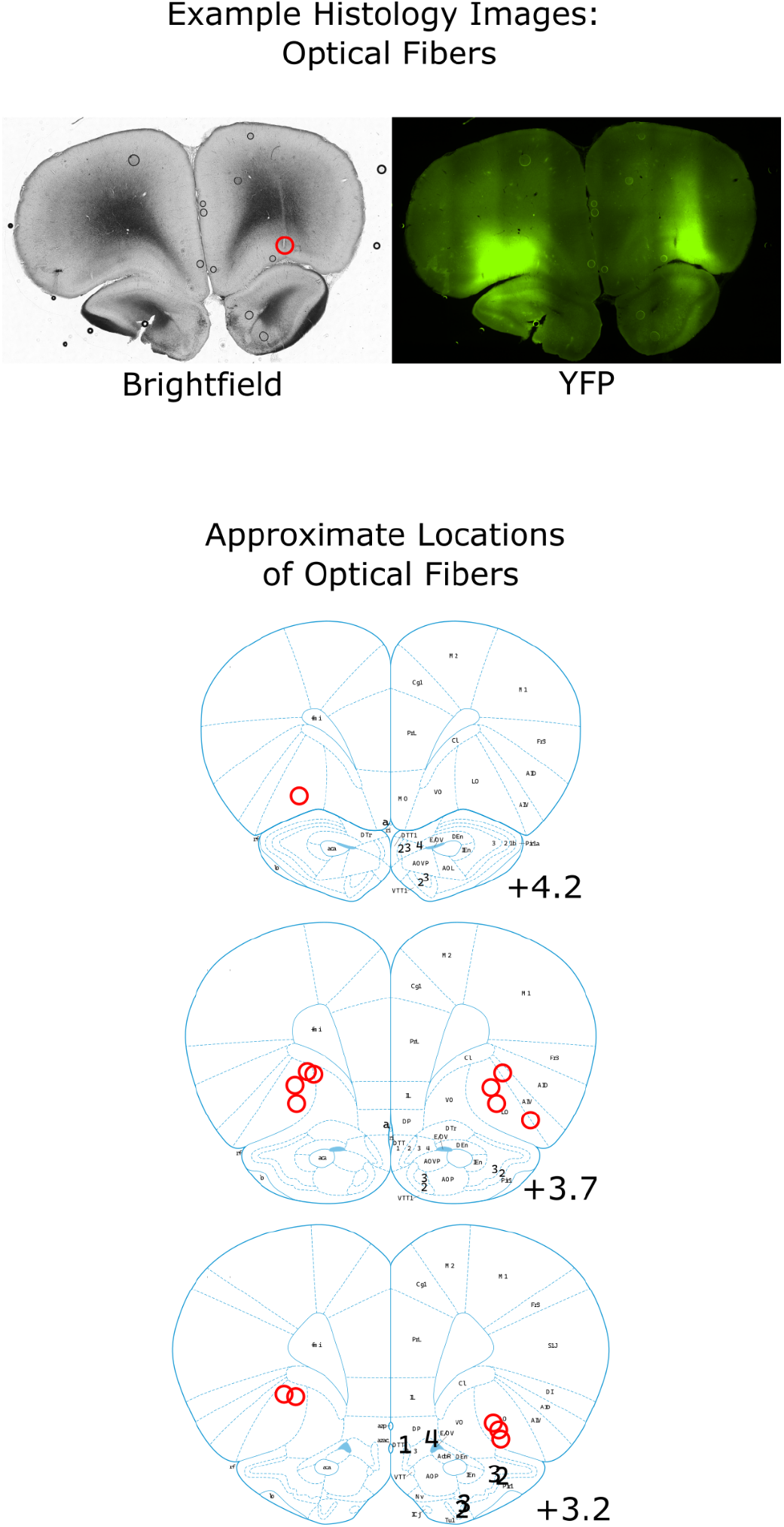
Locations of Optogenetics Implants. Example brightfield and YFP images of a coronal section taken from a rat implanted with optical fibers and infected with AAV-halorhodopsin. The location of the optical fiber on the right is visible as a scar (optical fiber on the left is visible in a different coronal section in this rat). The area of expression is visible in the YFP channel. Below: Estimates of the locations of all recording electrodes and fiber tips, obtained by comparing histology images to the reference atlas (Paxinos and Watson, 2006).

**Figure 5 - Supplement 2:**
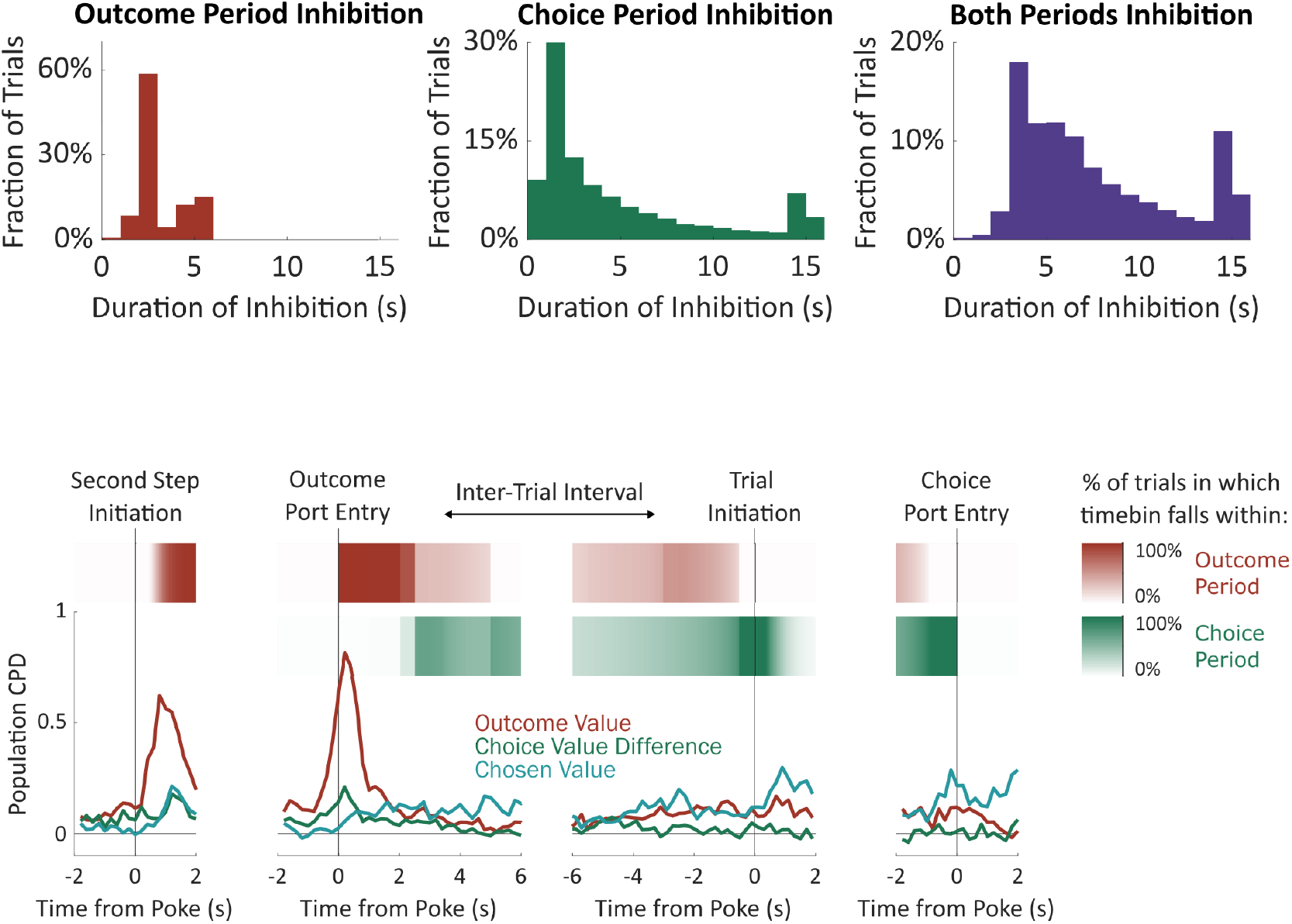
Periods of optogenetic inhibition. The “outcome period” on each trial was defined as beginning when the rat entered the outcome port, and ending either when the rat had left the outcome port for a continuous period of one second or more or after 2.5 seconds (unrewarded trials) or 5 seconds (rewarded trials). The “choice period” was defined as beginning at the end of the outcome period, and ending when the rat entered the choice port on the subsequent trial. “Both periods” inactivation began at entry into the outcome port and ended at entry into the choice port on the subsequent trial. For both “choice period” and “both periods” inactivation, an upper limit of 15 seconds was imposed. Trials on which inactivation ended due to this limit were not analyzed. **Above**: Histograms of the total duration of inactivation in each condition. **Below**: Relationship between the timebins used in electrophysiology analysis and the optogenetic inactivation periods. Line plot shows the population CPD for each of the three value-related regressors from the electrophysiology experiment (identical to Figure 4b). Red and green stripes show the probability of each timebin being in each of the optogenetic inactivation periods, computed using the optogenetics dataset.

**Figure 5 - Supplement 3:**
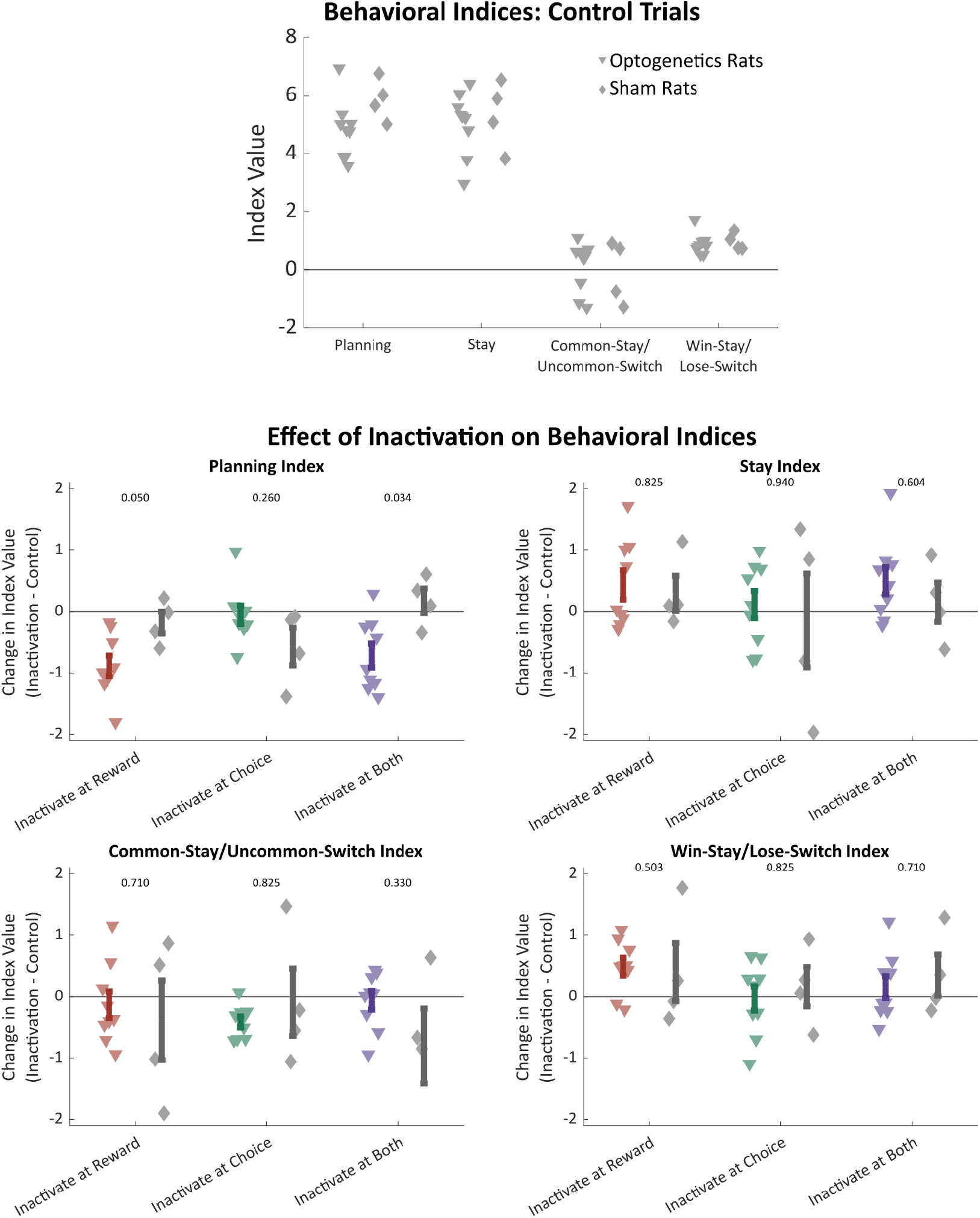
Effects of optogenetics on all behavioral indices. **Above**: Regression-based behavioral indices computed considering only trials that were not preceded by inactivation, shown separately for optogenetics rats (triangles) and sham optogenetics rats (diamonds). Behavior was similar in the two groups of rats. **Below**: Difference in behavioral indices between trials that were preceded by inactivation in each time period and those that were not preceded by inactivation. Only for the planning index were there significant differences between optogenetics and sham optogenetics rats (p-values shown from two-sample rank sum test).

**Figure 5 - Supplement 4:**
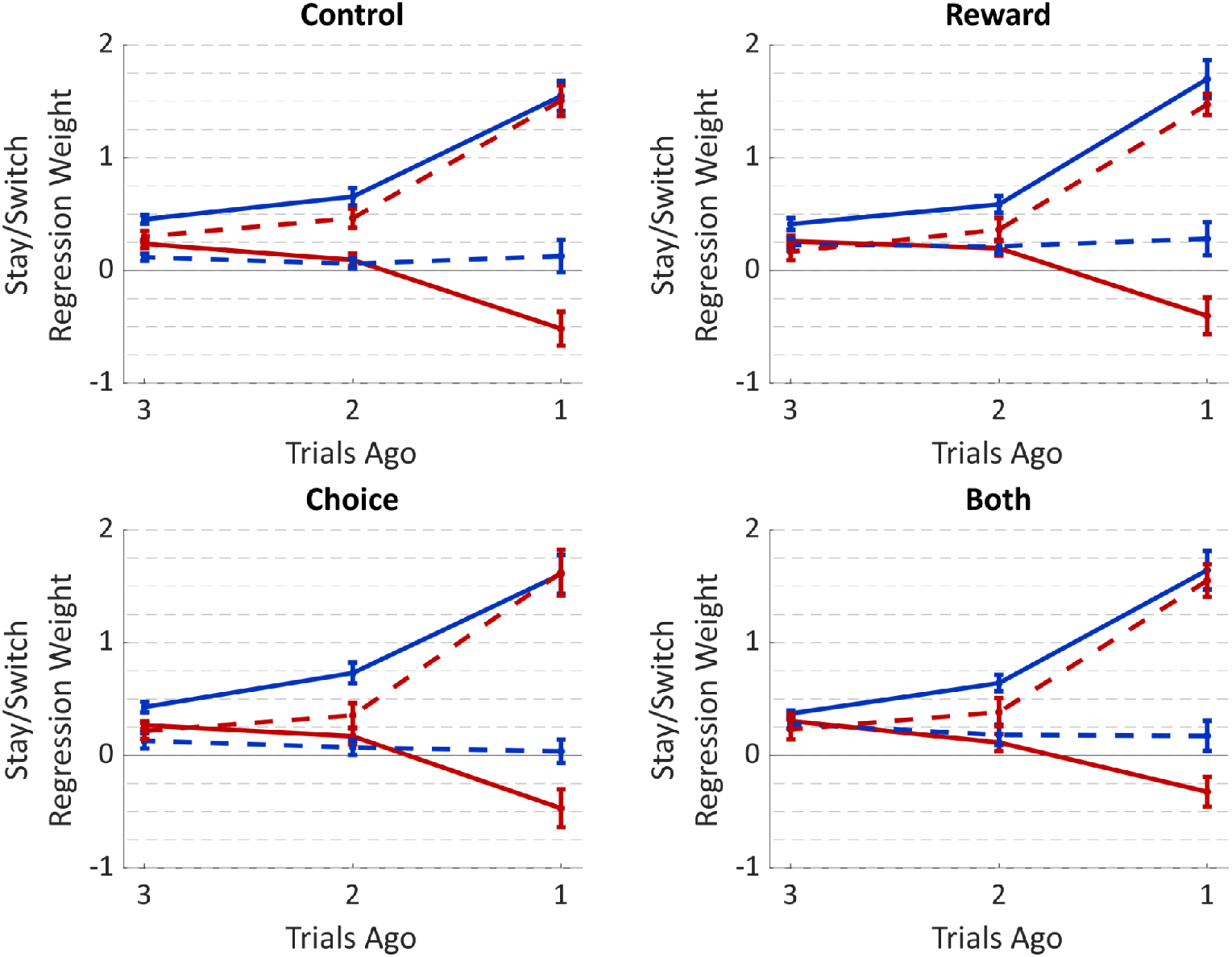
Average fit weights of the trial-history regression model to behavioral data from the optogenetics experiment. Weights were estimated separately for trials preceded by (**a**) no inhibition, (**b**) inhibition during the reward period, (**c**) inhibition during the choice period, or (**d**) inhibition during both periods. Weights for individual rats are shown in Figure 5-4

**Figure 5 - Supplement 5:**
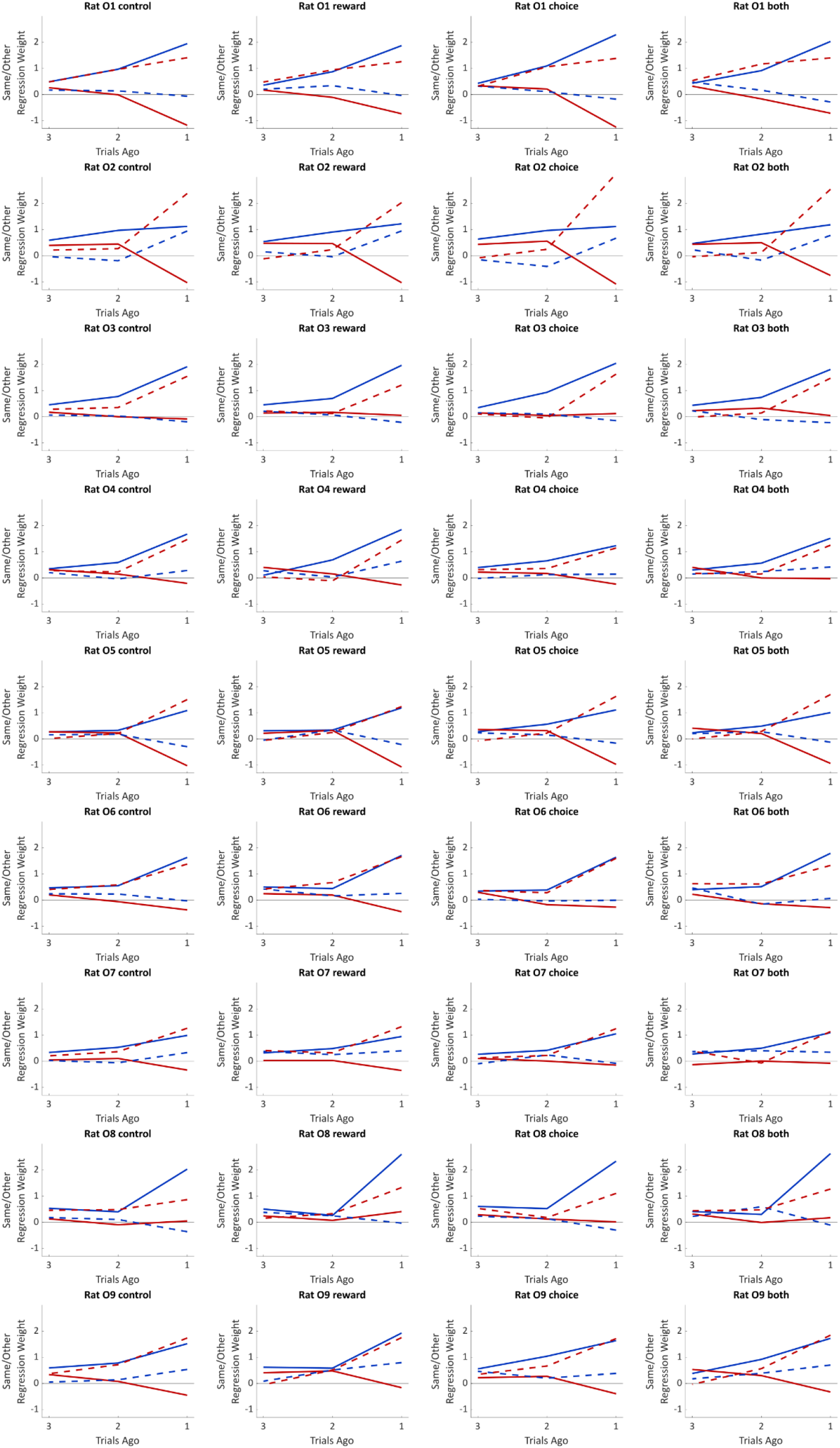
Fit weights of the trial history regression model for individual rats in the optogenetics experiment. Weights were estimated separately for trials preceded by no inhibition, inhibition during the reward period, inhibition during the choice period, or inhibition during both periods.

